# ESRP1-Mediated Alternative Splicing During Oocyte Development is Required for Mouse Fertility

**DOI:** 10.1101/2020.07.16.206425

**Authors:** Luping Yu, Huiru Zhang, Xuebing Guan, Dongdong Qin, Jian Zhou, Xin Wu

## Abstract

Alternative splicing (AS) contributes to gene diversification in cells, but the importance of AS during germline development remains largely undefined. Here, we interrupted pre-mRNA splicing events controlled by epithelial splicing regulatory protein 1 (ESRP1) and found that it induced female infertility in mice. Germline-specific knockout of *Esrp1* perturbed spindle organization, chromosome alignment, and metaphase-to-anaphase transformation in oocytes. The first polar body extrusion (PBE) was blocked during oocyte meiosis and was found to be due to abnormal activation of spindle assembly checkpoint (SAC) and insufficiency of anaphase-promoting complex/cyclosome (APC/C) in *Esrp1*-knockout oocytes. *Esrp1*-knockout in oocytes hampered follicular development and ovulation; eventually, premature ovarian failure (POF) occurred in six-month-old *Esrp1*-knockout mouse. Using single-cell RNA sequencing analysis, 528 aberrant AS events of maternal mRNA transcripts were revealed and were preferentially associated with microtubule cytoskeletal organization in *Esrp1*-knockout oocytes. Notably, we found that loss of ESRP1 disturbed a comprehensive set of gene-splicing sites—including those within *Trb53bp1, Rac1, Bora, Kif2c, Kif23, Ndel1, Kif3a, Cenpa*, and *Lsm14b*—that ultimately caused abnormal spindle organization. Taken together, our findings provide the first report elucidating the AS program of maternal mRNA transcripts, mediated by the splicing factor, ESRP1, that is required for oocyte meiosis and female fertility in mice.

## Introduction

Alternative splicing (AS) of pre-mRNA is a key step that gives rise to functionally distinct proteins from a single gene, according to the developmental or physiological state of cells in multicellular organisms. Variability in splicing patterns is a major source of protein diversity from the genome (Black 2000). Nearly 95%, 60%, and 25% of genes in humans, *Drosophila melanogaster*, and *Caenorhabditis elegans*, respectively (Wang et al. 2008; Graveley et al. 2011; Ramani et al. 2011), undergo alternative splicing based on RNA sequencing. Thus, AS significantly expands the forms and functions of the genomes of organisms with limited numbers of genes (Nilsen and Graveley 2010). Aberrations in splicing patterns have been implicated in a number of different diseases; for example, abnormal splicing patterns of fibroblast growth factor receptor 2 (*Fgfr2*) result in inner-ear developmental defects and hearing loss (Rohacek et al. 2017). Additionally, aberrations in the splicing patterns of G-protein-coupled receptor 137 (*Gpr137*) impair epithelial cell integrity and contribute to intestinal pathogenesis (Mager et al. 2017). However, the involvement of AS regulation in germline cell development has not been well elucidated.

The mammalian oocyte represents a unique physiological case because it exhibits high transcriptional activity during growth, followed by transcriptional quiescence when it is stimulated to resume meiosis. In non-surrounded nucleolus (NSN)-type oocytes, chromatin does not surround the nucleolus, and gene transcription is globally active; in contrast, in surrounded nucleolus (SN)-type oocytes, chromatin is highly condensed and gathered around the nucleolus, and gene transcription is globally silenced (Kageyama et al. 2007). Oocytes rely on the storage of maternal transcripts to maintain cellular processes throughout meiotic maturation and cytoplasmic maturational events that provide competence for fertilization and embryogenesis. Thus, a specific post-transcriptional regulatory context has increased importance in oocytes and is essential to generate high-quality female gametes (Sha et al. 2019; Christou-Kent et al. 2020). At present, some evidence has suggested that alternative pre-mRNA splicing significantly contributes to accurate control of maternal transcripts (Do et al. 2018; Kasowitz et al. 2018). However, the landscape of pre-mRNA splicing within the stored RNA pool in oocytes has not yet been critically examined.

RNA-binding proteins (RBPs) control the fate of RNAs with one or more RNA-recognition motif (RRM) domains and accessory domains, which participate in many post-transcriptional processes, including the splicing of pre-mRNA, as well as ensuring the localization and stability of RNAs within the cell (Hentze et al. 2018). Epithelial splicing regulatory protein 1 (ESRP1) is preferentially known as an epithelial-specific RBP that is composed of three RRM domains and regulates AS by directly binding specific GU-rich sequence motifs in pre-mRNAs, and plays a role in the epithelial-cell-specific splicing program (Warzecha et al. 2009b; Warzecha et al. 2010). ESRP1 also acts to maintain an epithelial-cell state by preventing the switch from CD44 variant isoforms (CD44v) to the standard isoform (CD44s), which is critical for regulating the epithelial-to-mesenchymal transition (EMT) and the progression of breast cancer (Brown et al. 2011). ESRP1 has been demonstrated to play essential roles in mammalian development through AS to maintain epithelial-cell properties (Bebee et al. 2015). However, the physiological role of ESRP1 in the maternal transcriptome in germline development has not yet been investigated. In the present study, we found that ESRP1-mediated AS was required for spindle organization, chromosome congression, meiotic progression, follicular development, and ovulation in mouse oocytes. Loss of ESRP1 led to arrest of mouse meiotic progression, ovulation disorders, premature ovarian failure, and female infertility.

## Results

### *Esrp1* knockout leads to female infertility in mice

A recent study used RBP capture to identify a set of RBPs with uncharacterized roles in mammalian germline cells, including ESRP1 (Du et al. 2020). Here, we validated the presence of ESRP1 in adult mouse stomach, lung, and kidney tissues (Fig. 1A). ESRP1 was also detectable in prepubertal mouse testes, while highly enriched expression of ESRP1 was found in mouse ovaries and at all follicular stages (Figs. 1A and S1A). Because ESRP1 is required for mouse craniofacial development and postnatal viability (Bebee et al., 2015), we next generated a conditional inactivation of the *Esrp1* gene using the Cre/loxP system to investigate the functional roles of *Esrp1* during germline development. In our experiments, the targeting construct contained three loxP sites inserted into intron 6 and intron 9 of the *Esrp1* gene and one floxed hygromycin and thymidine kinase (HyTK) double-selection cassette. All three loxP sites were in the same orientation (Fig. S2A). Mice carrying the floxed allele (*Esrp1*^*fl/fl*^) were backcrossed with *Ddx4-Cre* mice (cat. #J006954, Jackson Laboratory) to produce mice with germ-cell-specific inactivation of *Esrp1* in primordial germ cells (PGCs) starting at embryonic day 12.5 (designated hereafter as *Esrp1*^*fl/Δ*^*/Ddx4-Cre*; mouse-mating strategies are illustrated in Fig. S2B). We first found that *Esrp1*-knockout male mice were grossly normal in terms of the morphology and size of the testes and epididymis, as compared to those of wild-type mice (Fig. 1B). Next, morphological analysis indicated that there were no apparent changes in spermatogenesis at any stages of the seminiferous tubules in eight-week-old *Esrp1*^*fl/Δ*^ */Ddx4-Cre* mouse testes (Fig. S1B). Furthermore, there was no significant decrease in male fertility (Fig. 1C); however, females were completely infertile (Fig. 1G), as indicated by the results of our breeding experiments from *Esrp1*-knockout mice. Collectively, these data suggest that loss of ESRP1 in germline cells led to female infertility, whereas such loss was dispensable for male fertility in mice.

**Figure 1.**
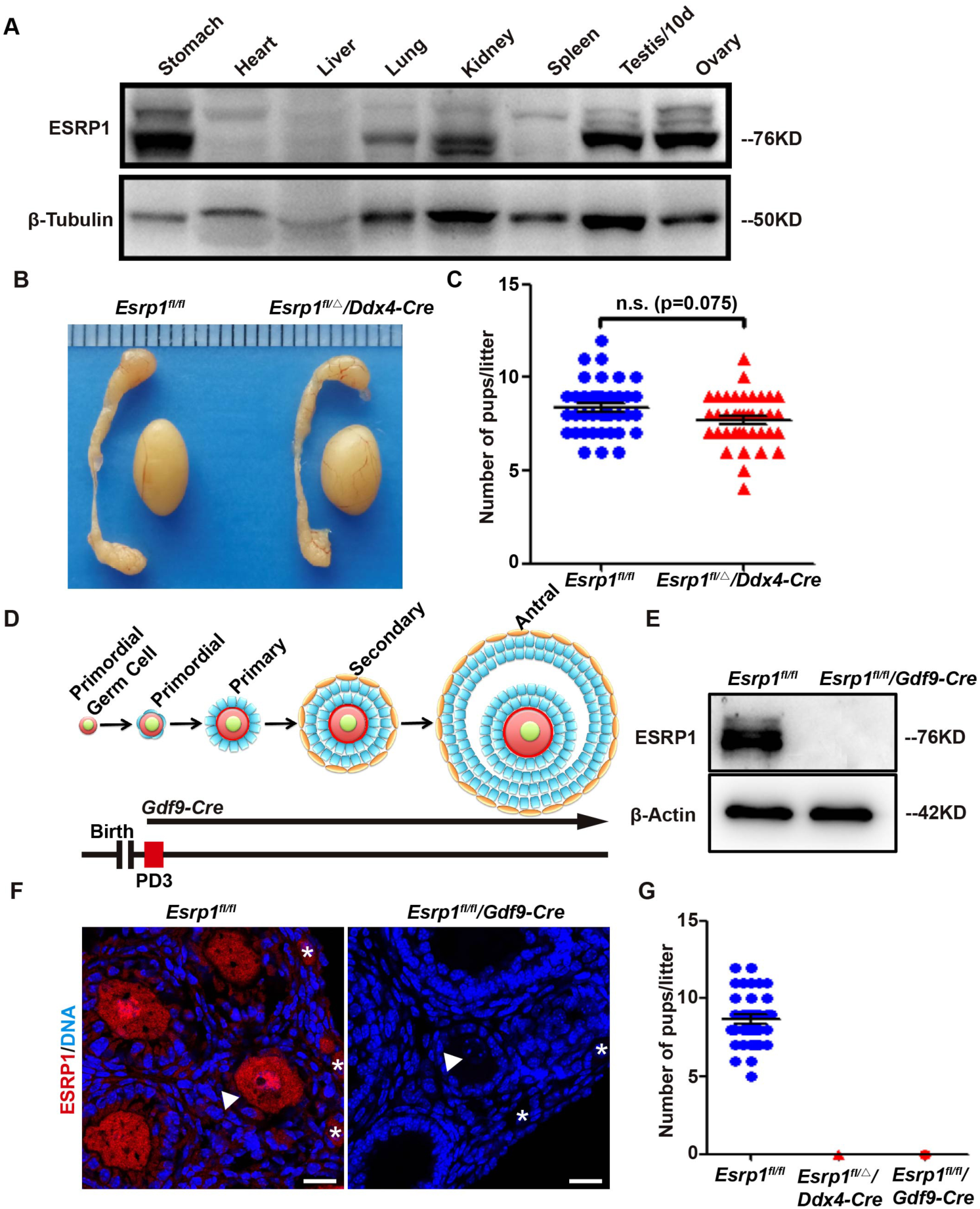
Germline deletion of ESRP1 leads to female infertility in mice. (A) ESRP1 expression in mouse tissues. Western-blot analyses showing the expression of ESRP1 protein in adult mouse stomach, lung, kidney, ovary, and prepubertal testicular tissues. β-tubulin served as a loading control. (B) Morphology of eight-week-old testes and epididymis from *Esrp1*^*fl/fl*^ and *Esrp1*^*fl/Δ*^*/Ddx4-Cre* mice. (C) Fertility analysis of *Esrp1*^*fl/fl*^ and *Esrp1*^*fl/Δ*^*/Ddx4-Cre* males. (D) Schematic diagram of the developmental time point at which *Gdf9-Cre* was expressed. (E) Western-blot analysis showing the expression of ESRP1 protein in GV oocytes from the ovaries of *Esrp1*^*fl/fl*^ and *Esrp1*^*fl/fl*^*/Gdf9-Cre* mice. (F) Immunofluorescent staining of ESRP1 (red) in PD21 mouse ovaries. DNA was stained by DAPI (blue). Asterisks indicate oocytes in primordial follicles and white arrowheads point to growing oocytes. Scale bars, 20 µm. (G) Fertility analysis of the indicated genotypes mice; *Esrp1*^*fl/Δ*^*/Ddx4-Cre* and *Esrp1*^*fl/fl*^ */Gdf9-Cre* female were infertile.

### Follicular developmental defects and POF in *Esrp1*-knockout mouse

In order to elucidate the requirement of postnatally expressed ESRP1 for female fertility, we bred *Esrp1*^*fl/fl*^ mice with male transgenic (Tg) mice [Tg(*Gdf9*-icre)5092Coo, cat. #J011062, Jackson Laboratory] to inactivate the *Esrp1* gene in oocytes starting at the primordial follicle at postnatal day 3 (PD3) (designated hereafter as *Esrp1*^*fl/fl*^*/Gdf9-Cre*, Fig. 1D) (Lan et al. 2004), following our previous breeding strategies (Fig. S2B). ESRP1 protein was not detectable in *Esrp1* mutant oocytes from *Esrp1*^*fl/fl*^*/Gdf9-Cre* mouse ovaries (Fig. 1E), and immunofluorescent analysis showed that ESRP1 was absent in primordial oocytes of *Esrp1*^*fl/fl*^*/Gdf9-Cre* ovaries and at all stages of subsequent oocytes (Fig. 1F). As we expected, no pregnancies in female mice and no progeny were observed when *Esrp1*^*fl/fl*^*/Gdf9-Cre* female mice were bred with wild-type males, as comparing to results in *Esrp1*^*fl/fl*^ female mice that produced approximately eight to nine pups per litter and approximately six litters per female during six-month breeding periods (Fig. 1G).

Next, we examined the gross morphology of ovaries at different ages and analyzed the composition of follicles according to the classifications of primordial, primary, early-secondary, later-secondary, antral follicles, and the corpus luteum. Follicle development was quantified to explore the reproductive function of female mice, and we found that *Esrp1*^*fl/fl*^*/Gdf9-Cre* ovaries were smaller in size compared to *Esrp1*^*fl/fl*^ ovaries from PD28 (Fig. 2A, D, G, J). The total number of follicles in *Esrp1*^*fl/fl*^*/Gdf9-Cre* ovaries was significantly lower compared to those in *Esrp1*^*fl/fl*^ mice at all stages (Fig. 2O). At PD28, there were fewer later-secondary and antral follicles (Fig. 3C’); from PD60, follicles at all stages were dramatically reduced due to follicle atresia (Fig. 2F’, I’, O); at PD180, nearly all follicles were absent except for a few primary, secondary, and antral follicles that were found in *Esrp1*^*fl/fl*^*/Gdf9-Cre* mouse ovaries (Fig. 2K, L, L’). Different from control ovaries, the layers of cumulus cells and mural granulosa cells (MGC) were significantly thinner at later-secondary and antral follicles in *Esrp1-*knockout ovaries (Figs. 2B, C, E, F, H, I, arrows and inserts), indicating that ESRP1 in oocyte was required for follicular development but may not be necessary for the transition of primordial oocytes to the activated growing oocyte stage. Notably, the corpus luteum was significantly less in *Esrp1-*knockout ovaries compared to that in *Esrp1*^*fl/fl*^ ovaries from PD28 to PD180 (Fig. 2C’, F’, I’, L’), suggesting that ovulation was impaired in *Esrp1-*knockout mice. We also administered intraperitoneal injections of pregnant mare serum gonadotropin (PMSG) and human chorionic gonadotropin (hCG) at 44–46 h after PMSG treatment to induce ovulation in six-week-old mice. In contrast to results in *Esrp1*^*fl/fl*^ oocytes, histological analysis of cumulus-oocyte complexes (COCs) ovulated in mouse oviducts showed that *Esrp1-*knockout oocytes did not extrude the first polar body (PB1) (Fig. 2M, N). Therefore, oocytes in *Esrp1*^*fl/fl*^*/Gdf9-Cre* mice lacked the competence to undergo meiotic maturation. These results suggest that ESRP1 was required for follicular development, ovulation and oocyte maturation in mice.

**Figure 2.**
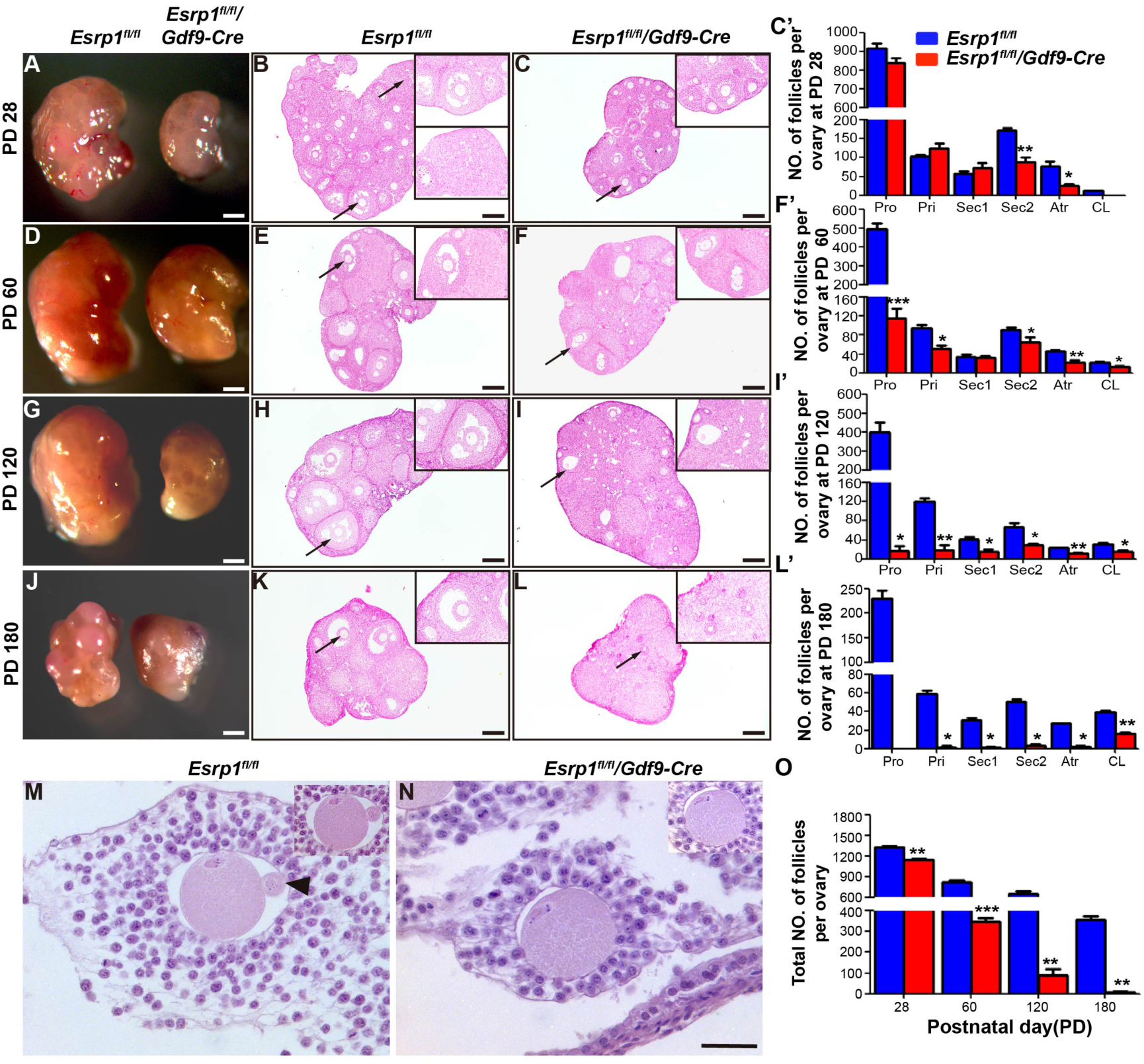
ESRP1 is essential for follicular development and oocyte growth. (A) Morphology of *Esrp1*^*fl/fl*^ and *Esrp1*^*fl/fl*^*/Gdf9-Cre* mouse ovaries at PD28. Scale bars, 500 µm. (B–C) Hematoxylin/eosin (HE) staining of paraffin slides of ovaries. Arrows and inserts show the antral follicles and corpus luteum at PD28 *Esrp1*^*fl/fl*^ mice ovaries (B) and antral follicles at PD28 *Esrp1*^*fl/fl*^ */Gdf9-Cre* ovaries (C). Scale bars, 100 µm. (C’) Statistical analysis of the numbers of follicles in the ovaries of *Esrp1*^*fl/fl*^ (B) and *Esrp1*^*fl/fl*^*/Gdf9-Cre* (C) mice (*p< 0.05, **p < 0.01). (D) Morphology of *Esrp1*^*fl/fl*^ and *Esrp1*^*fl/fl*^*/Gdf9-Cre* mouse ovaries at PD60. Scale bars, 500 µm. (E–F) HE staining showing that the layers of cumulus cells and mural granulosa cells (MGC) of antral follicles at PD60 *Esrp1*^*fl/fl*^ */Gdf9-Cre* mice ovaries (F, arrow and insert) were significantly thinner compared to those in *Esrp1*^*fl/fl*^ mice (E, arrow and insert). Scale bars, 100 µm. (F’) Statistical analysis of the numbers of follicles in the ovaries of *Esrp1*^*fl/fl*^ (E) and *Esrp1*^*fl/fl*^*/Gdf9-Cre* (F) mice (*p< 0.05, **p < 0.01, ***p < 0.001). (G) Morphology of *Esrp1*^*fl/fl*^ and *Esrp1*^*fl/fl*^*/Gdf9-Cre* mouse ovaries at PD120. Scale bars, 500 µm. (H–I) HE staining showing that the layers of MGC of antral follicles at PD120 *Esrp1*^*fl/fl*^ */Gdf9-Cre* mice (I, arrow and insert) were significantly thinner compared to those in *Esrp1*^*fl/fl*^ mice (H, arrow and insert). Scale bars, 100 µm. (I’) Statistical analysis of the numbers of follicles in the ovaries of *Esrp1*^*fl/fl*^ (E) and *Esrp1*^*fl/fl*^ */Gdf9-Cre* (F) mice (*p< 0.05, **p < 0.01). (J) Morphology of *Esrp1*^*fl/fl*^ and *Esrp1*^*fl/fl*^*/Gdf9-Cre* mouse ovaries at PD180. Scale bars, 500 µm. (K–L) HE staining showing that nearly all follicles were absent at PD180 in *Esrp1*^*fl/fl*^*/Gdf9-Cre* mice (L, arrow and insert) compared to those in *Esrp1*^*fl/fl*^ mice (K, arrow and insert). Scale bars, 100 µm. (L’) Statistical analysis of the numbers of follicles in the ovaries of *Esrp1*^*fl/fl*^ (K) and *Esrp1*^*fl/fl*^*/Gdf9-Cre* (L) mice (*p< 0.05, **p < 0.01). (M–N) Histology of cumulus-oocyte complexes ovulated in oviducts in *Esrp1*^*fl/fl*^ (M) and *Esrp1*^*fl/fl*^*/Gdf9-Cre* (N) mice at 16 h after hCG injections. Black arrowheads indicate the first polar body. Scale bars, 100 µm. (O) The total numbers of follicles in *Esrp1*^*fl/fl*^ and *Esrp1*^*fl/fl*^ */Gdf9-Cre* mice at different ages (*p< 0.05, **p < 0.01, ***p < 0.001). Abbreviations: Pro: primordial follicles; Pri: primary follicles; Sec1: early-secondary; Sec2: later-secondary; Atr: antral follicles; CL: corpus luteum.

**Figure 3.**
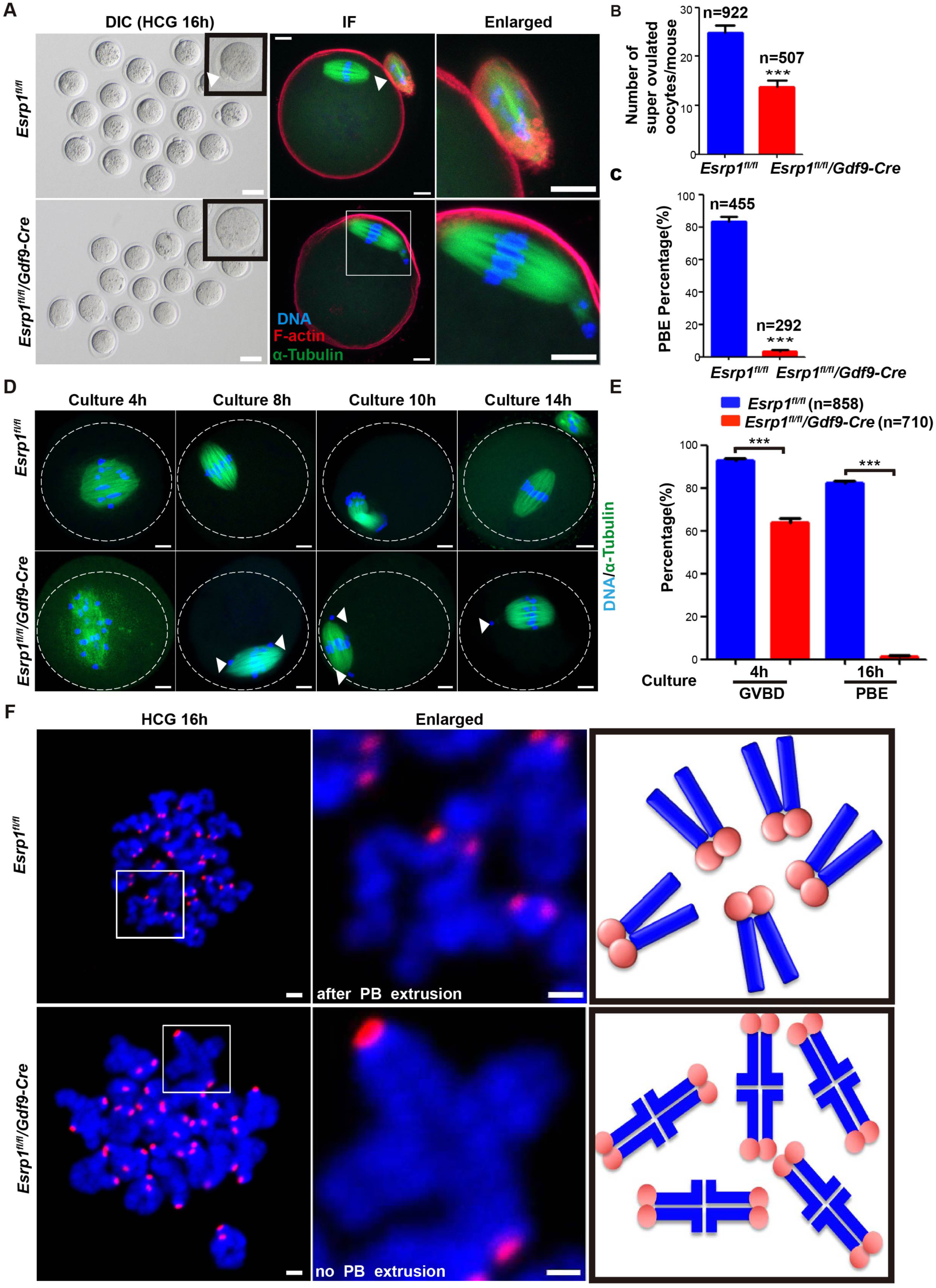
ESRP1 is required for the metaphase-to-anaphase-I transition in oocytes. (A) Differential interference contrast (DIC) images of ovulated oocytes from *Esrp1*^*fl/fl*^ and *Esrp1*^*fl/fl*^*/Gdf9-Cre* mice at 16 h after hCG injections (left). Scale bars, 100 µm. Immunofluorescent staining with α-tubulin (green), F-actin (red), and DAPI (blue) in ovulated oocytes from *Esrp1*^*fl/fl*^ and *Esrp1*^*fl/fl*^*/Gdf9-Cre* mice (right). Scale bars, 10 µm. (B) Numbers of oocytes ovulated by *Esrp1*^*fl/fl*^ mice (24.82 ± 1.47) and *Esrp1*^*fl/fl*^*/Gdf9-Cre* mice (13.70 ± 1.40) (***P < 0.001). n: number of used oocytes. (C) The rates of MII oocytes with the PB1 from superovulated *Esrp1*^*fl/fl*^ mice (83.54% ± 2.79%) and *Esrp1*^*fl/fl*^*/Gdf9-Cre* mice (3.10% ± 1.23%) (***P < 0.001). n: number of used oocytes. (D) Immunofluorescent of spindle stained with α-tubulin (green) and chromosomes with DAPI (blue) in oocytes of *Esrp1*^*fl/fl*^ mice (n=60) and *Esrp1*^*fl/fl*^ */Gdf9-Cre* mice (n=58) at the indicated time points. Dashed lines indicate oocyte outlines, and white arrows indicate lagging chromosomes. Scale bar, 10 mm. (E) GVBD rates of cultured oocytes that were collected at the GV stage from *Esrp1*^*fl/fl*^ mice (92.86% ± 0.89%) and *Esrp1*^*fl/fl*^*/Gdf9-Cre* mice (64.07% ± 1.56%) (***P < 0.001). PBE rates of *Esrp1*^*fl/fl*^ mice (82.09% ±1.19%) and *Esrp1*^*fl/fl*^*/Gdf9-Cre* mice (1.10% ± 0.53%) (***P < 0.001). n: number of used oocytes. (F) Chromosome spreads of ovulated oocytes from superovulated *Esrp1*^*fl/fl*^ and *Esrp1*^*fl/fl*^ */Gdf9-Cre* mice at 16 h after hCG injections. Kinetochores were stained with CREST (red), and chromosomes were stained with DAPI (blue). The illustrations show typical chromosomes that were observed. Scale bar, 1 µm.

### Loss of ESRP1 induces oocyte meiotic arrest at metaphase I

Next, we looked for possible defects during oocyte development in *Esrp1*^*fl/fl*^*/Gdf9-Cre* female mice. Using superovulation experiments, we first found an average of approximately twenty-five ovulated oocytes in *Esrp1*^*fl/fl*^ females after 16 h of hCG treatments (Fig. 3B); however, the ovulation rate in *Esrp1*^*fl/fl*^*/Gdf9-Cre* females was significantly reduced to an average of approximately fourteen oocytes (Fig. 3B). In contrast to normal oocytes ovulated from *Esrp1*^*fl/fl*^ ovaries, displaying the PB1 and a typical MII spindle (Fig. 3A, upper panel, differential interference contrast and immunofluorescence staining, white arrow indicated) in mature MII-stage oocytes, almost all of the ovulated oocytes from *Esrp1*^*fl/fl*^*/Gdf9-Cre* ovaries had undergone germinal vesicle breakdown (GVBD) with a spindle that had migrated to the cortex but lacked the PB1 (Fig. 3A, low panel). The polar body extrusion (PBE) percentage in *Esrp1*^*fl/fl*^*/Gdf9-Cre* females was significantly decreased compared to that in *Esrp1*^*fl/fl*^ females (Fig. 3C). Next, we explored whether the defects in oocyte development found *in vivo* could be reproduced through *in-vitro* maturation (IVM) experiments. We first found that *Esrp1-*knockout oocytes had successive occurrences of GVBD within 4 h, but with many lagged chromosomes and the percentage of GVBD oocytes was reduced after 4 h (Fig. 3E). Importantly, only 6 in total 710 oocytes were found PBE after 16 h during IVM in *Esrp1*^*fl/fl*^*/Gdf9-Cre* females, while 82.09% *Esrp1*^*fl/fl*^ oocytes had entered into the metaphase-II stage, as characterized by visible PB1 and univalent sister chromatids (Figs. 3D, E, S3A, B). None of the *Esrp1*^*fl/fl*^*/Gdf9-Cre*-ovulated oocytes underwent successful fertilization and developed into blastocysts after *in-vitro* fertilization (IVF) with normal sperm (Fig. S3C). Using chromosome-spread assay (Hodges and Hunt 2002), we further analyzed chromosome morphology in ovulated oocytes from *Esrp1*^*fl/fl*^/Gdf9-Cre mice. Oocytes from *Esrp1*^*fl/fl*^ mice displayed the typical monovalent MII-stage chromosome array (pairs of sister chromatids juxtaposed near their centromeres), whereas *Esrp1*^*fl/fl*^*/Gdf9-Cre* oocytes had intact bivalents with no homologue separation (Fig. 3F). These observations strongly indicate that *Esrp1-*knockout oocytes were arrested at metaphase I with inseparable homologous chromosomes.

### ESRP1 deficiency hampers spindle/chromosome organization and K-MT attachments

Next, we investigated whether defects occurred in spindle/chromosome organization therefore contributed to metaphase-I arrest in *Esrp1*^*fl/fl*^*/Gdf9-Cre* oocytes. We found that the incidence of spindle defects and chromosome misalignments were significantly higher in *Esrp1*^*fl/fl*^*/Gdf9-Cre* ovulated oocytes than in *Esrp1*^*fl/fl*^ oocytes (94.05% ± 1.62% vs. 15.02% ± 1.75%; spindles in *Esrp1-*knockout oocytes retained the barrel-shape characteristic of metaphase I even at ovulated oocytes considered to have abnormal spindle/chromosome organization, Fig. 4A, B). Compared to *Esrp1*^*fl/fl*^ oocytes (typical barrel-shaped MII spindles with organized chromosomes at the equator), a large portion of *Esrp1-*knockout oocytes (48.86%) contained abnormal spindle/chromosome organization with apparent monopolar spindle attachments (either single or multiple chromosomes); 39.77% of oocytes with MI arrest spindles; 9.09% of oocytes exhibited catastrophic spindle disorganization and severely misaligned chromosomes and 2.27% of oocytes had multiple spindles (Fig. 4A, C). These results indicate that ESRP1 was required for spindle organization and chromosome alignment during meiosis in mice.

**Figure 4.**
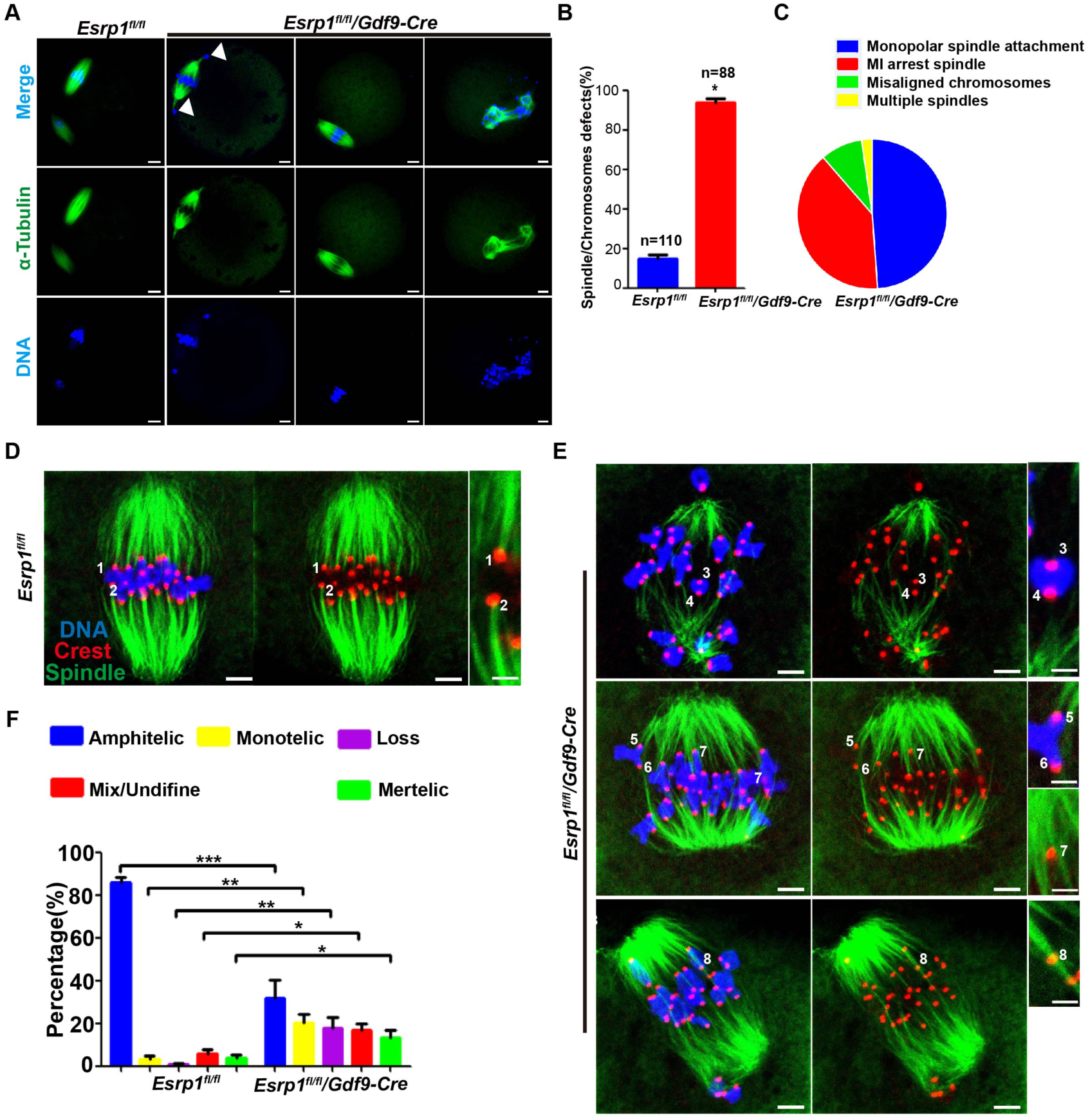
*Esrp1* deletion leads to spindle/chromosome disorganization and impaired K-MT attachments during oocyte meiosis. (A) Immunofluorescent staining with α-tubulin to visualize spindles (green) and co-stained with DAPI to visualize chromosomes (blue) in ovulated oocytes of *Esrp1*^*fl/fl*^ and *Esrp1*^*fl/fl*^*/Gdf9-Cre* mice. Left, *Esrp1*^*fl/fl*^ ovulated oocytes presented the PB1, a typical barrel-shaped spindle and well-aligned chromosomes on the metaphase plate; Right, *Esrp1*^*fl/fl*^*/Gdf9-Cre* ovulated oocytes lacked the PB1, and three examples illustrate apparent monopolar spindle attachments (arrows), metaphase-I arrest spindles, and catastrophic spindle disorganization that were frequently observed in *Esrp1*-knockout oocytes. Representative confocal sections are shown. Scale bar, 10 µm. (B) The incidence of abnormal spindles and misaligned chromosomes in *Esrp1*^*fl/fl*^ and *Esrp1*^*fl/fl*^*/Gdf9-Cre* oocytes (*P < 0.05). n: number of used oocytes. (C) The frequency distribution of *Esrp1*^*fl/fl*^*/Gdf9-Cre* oocyte phenotypes, within which one oocyte may fall into more than one phenotypic class. (D–E) Control and *Esrp1*-knockout oocytes at MI stage staining with α-tubulin to visualize spindles (green), CREST to detect kinetochore (red), and co-stained with DAPI to visualize chromosomes (blue). (D) Representative confocal image showing the amphitelic attachment in control oocyte (chromosomes 1 and 2). Scale bar, 5 µm or 2 µm. (E) Representative confocal images showing the lost attachment (chromosomes 3 and 4), monotelic attachment (chromosomes 5 and 6), mix/undefined attachment (chromosome 7), as well as merotelic attachment (chromosome 8) in *Esrp1*-knockout oocytes. Scale bar, 5 µm or 2 µm. (F) Quantitative analysis of K-MT attachments in control and *Esrp1*-knockout oocytes. Kinetochores in regions where fibers were not easily visualized were not included in the analysis. 12 control oocytes and 12 *Esrp1*-knockout oocytes were examined respectively (*p< 0.05, **p < 0.01, ***p < 0.001).

On meiotic entry, dynamic microtubules form a bipolar spindle, which is responsible for capturing and congressing chromosomes. These events require proper attachment of kinetochores to microtubules emanating from opposite spindle poles(Touati et al. 2015). We investigated whether kinetochore-microtubule (K-MT) interactions were compromised resulting in defects contributed to spindle/chromosome organization in *Esrp1-*knockout oocytes. To do this, metaphase I oocytes were labeled with α-tubulin to visualize spindle and CREST to detect kinetochores, and co-stained with DAPI for chromosomes (Fig. 4D, E). We found that the majority K-MT pattern in normal oocytes is amphitelic attachment (the kinetochore of one chromosome is connected to the spindle pole and the kinetochore of the other chromosome is connected to the opposite spindle pole; Fig. 4 D, F, chromosomes labeled 1 and 2). By contrast, the proportion of amphitelic K-MT attachment was significantly reduced and the frequency of misattachments was significantly increased in *Esrp1-*knockout oocytes relative to control (Fig. 4F), including lost attachment (kinetochore attached to neither pole; Fig.4E, chromosomes labeled 3 and 4), monotelic attachment (Fig. 4E, chromosomes labeled 5 and 6), mix/undefined attachment (Fig. 4E, chromosome labeled 7), as well as merotelic attachment (one kinetochore attached to both poles; Fig. 4E, chromosomes labeled 8). The results indicate that the erroneous K-MT attachments could result in chromosome alignment failure observed in *Esrp1-*knockout oocytes. Together, ESRP1 is essential for K-MT interactions during oocyte meiotic maturation.

### ESRP1 deficiency activates the SAC and induces insufficient APC/C activity

The spindle assembly checkpoint (SAC) ensures correct chromosome alignment during meiosis (Overlack et al. 2015). Since *Esrp1*^*fl/fl*^ */Gdf9-Cre* oocytes failed to complete meiosis-I division and exhibited severe spindle-organization defects and abnormal chromosome alignments, we next evaluated the activity of the SAC in *Esrp1-*knockout oocytes. We analyzed the levels of BubR1, a principal component of the SAC in oocytes. Oocytes collected from *Esrp1*^*fl/fl*^ and *Esrp1*^*fl/fl*^*/Gdf9-Cre* mice were used to evaluate the SAC activity at 4 h and 6 h after GVBD (the time points presenting oocytes at premetaphase-I (pre-MI) and metaphase-I (MI) stages, respectively). In *Esrp1*^*fl/fl*^ oocytes, BubR1 was localized to kinetochores during pre-MI, and then degenerated during MI when kinetochores had become properly attached to microtubules (Fig. 5A, B). In contrast, the BubR1 signal on kinetochores was dramatically increased in *Esrp1-*knockout MI oocytes (Fig. 5A, B), suggesting that SAC was activated.

**Figure 5.**
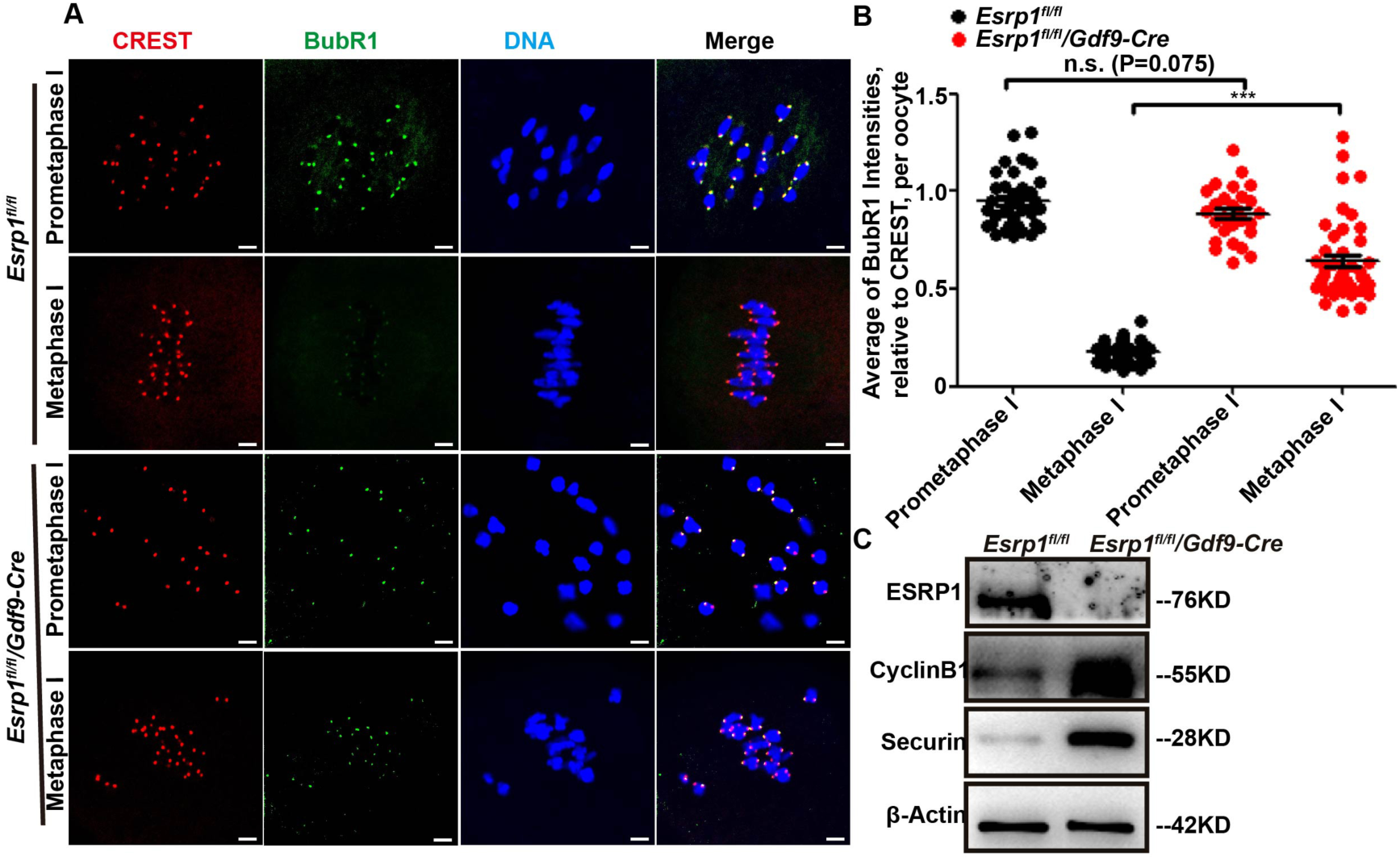
*Esrp1* knockdown activates SAC and induces insufficient APC/C activity. (A) Chromosome spreads were prepared 4 h (prometaphase-I) and 6 h (metaphase-I) after GVBD in oocytes of *Esrp1*^*fl/fl*^ and *Esrp1*^*fl/fl*^*/Gdf9-Cre* mice, then stained with CREST (red), BubR1 (green) and DAPI (blue). Scale bar, 5 µm. (B) Quantification of BubR1 signal intensity relative to CREST; each point is the relative intensity across centromeres. Data from prometaphase-I *Esrp1*^*fl/fl*^ oocytes (0.95 ± 0.02) and *Esrp1*^*fl/fl*^ */Gdf9-Cre* oocytes (0.88 ± 0.03) are shown. Metaphase-I *Esrp1*^*fl/fl*^ oocytes (0.18 ± 0.01) and *Esrp1*^*fl/fl*^ */Gdf9-Cre* oocytes (0.64 ± 0.03) are shown (***P < 0.001). (C) Western-blot analysis of Cyclin B1 and Securin in oocytes collected after 10 h in culture from *Esrp1*^*fl/fl*^ and *Esrp1*^*fl/fl*^*/Gdf9-Cre* mice. β-actin served as a loading control. Experiments were repeated at least three times; representative images are shown.

Complete activation of anaphase-promoting complex/cyclosome (APC/C) requires fulfillment of the SAC (Holt et al. 2013; Touati and Wassmann 2016). Since we found that the SAC was abnormally activated in *Esrp1-*knockout oocytes, we next asked whether MI arrest in *Esrp1-*knockout oocytes was due to insufficient APC/C activity. We collected 10-h cultured oocytes from *Esrp1*^*fl/fl*^ and *Esrp1*^*fl/fl*^*/Gdf9-Cre* females and assessed the protein levels of cyclin B1 and securin, two substrates of APC/C, to evaluate APC/C activity of oocytes during the metaphase-to-anaphase transition (Thornton and Toczyski 2003). In *Esrp1*^*fl/fl*^ oocytes, both proteins were nearly undetectable; however, in *Esrp1-*knockout oocytes, both proteins were present during the metaphase-to-anaphase transition (Fig. 5C), suggesting that there was no effective APC/C activity for cyclin B1 and securin degradation, rendering *Esrp1-*knockout oocytes unable to enter into anaphase. Therefore, the SAC surveillance mechanism and APC/C activity might represent major pathways mediating the effects of loss of ESRP1 on meiotic progression in oocytes.

### Inhibition of SAC activity rescues defects of metaphase-I arrest induced by ESRP1 deficiency

Since we found that conditional knockout of *Esrp1* led to SAC activity during MI and led to insufficient APC/C activity, we next investigated whether direct interference of SAC activity during MI could rescue the phenotype of *Esrp1-*knockout oocytes. We treated *Esrp1-*knockout oocytes with reversine (Fig. 6A), which inhibits the SAC kinase, MPS1 (Santaguida et al. 2010). Reversine accelerate meiosis I progression in *Esrp1*^*fl/fl*^ oocytes and the PBE rate is adjusted to 100% 2h after reversine added, as previously shown (Sha et al. 2018; Zhang et al. 2020) (Fig. 6B, C, D). Surprisingly, reversine partially rescued metaphase I arrest caused by *Esrp1* deficiency and led to the PB1 extrusion (Fig. 6B, C, D). The PB1 extrusion was confirmed by immunofluorescence assay (Fig. 6E). The recovered PB1 extrusion, characterized by homologous chromosome separation and disappearance of bivalent chromosomes, was confirmed by chromosome-spread assay (Fig. 6F). Therefore, we conclude that inhibition of SAC activity rescued metaphase-I arrest caused by *Esrp1* deficiency.

**Figure 6.**
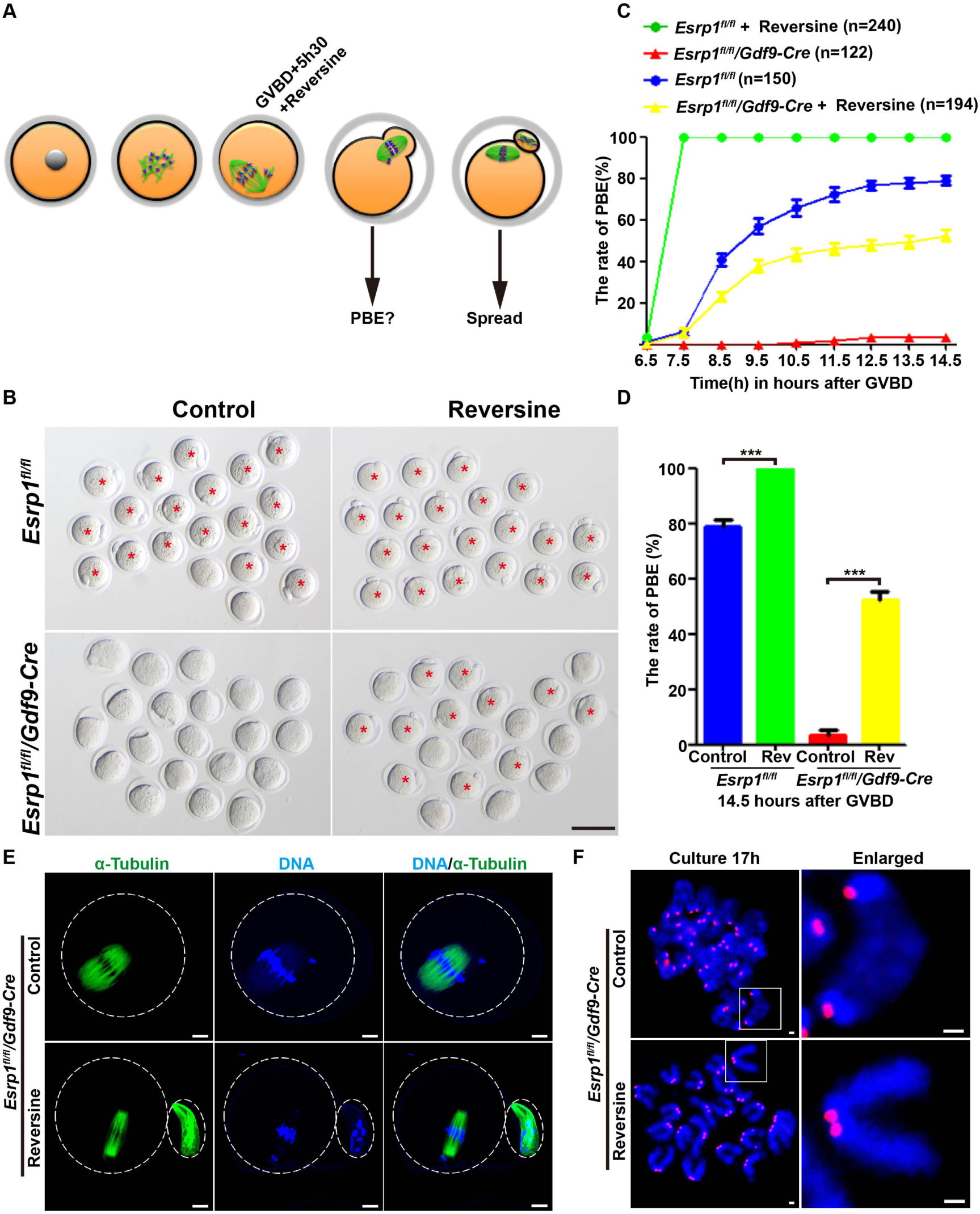
Metaphase-I arrest in *Esrp1*-knockout oocytes is rescued via inhibition of SAC activity. (A) Oocytes were treated with reversine (0.5 µM) at the indicated time to override a potential SAC arrest. (B) DIC imaging of *Esrp1*^*fl/fl*^ and *Esrp1*-knockout oocytes treated with reversine shows that they completed the first meiosis, as determined by PBE cultured for 16.5 h. The asterisk indicates PB1. Scale bar, 100 µm. At least three independent experiments involving a total of at least six mice were performed. (C) Time of PBE of *Esrp1*^*fl/fl*^ and *Esrp1*^*fl/fl*^*/Gdf9-Cre* oocytes. Reversine was added at GVBD+5.5 h, where indicated. n: number of used oocytes. (D) Statistical analysis of the PBE rates of oocytes shown in (B). Data from *Esrp1*^*fl/fl*^ oocytes (control: 78.96% ± 2.31%) and (reversine: 100%) are shown. Data from *Esrp1*^*fl/fl*^ */Gdf9-Cre* oocytes (control: 3.49% ± 1.78%) and (reversine: 52.58%± 2.51%) are shown (***P < 0.001). Abbreviations: rev: reversine. (E) Immunofluorescent staining with α-tubulin to visualize spindles (green) and co-stained with DAPI to visualize chromosomes (blue) show *Esrp1*-knockout oocytes had extruded PB1 after reversine treatment. Scale bar, 10 µm. (F) Oocytes were spread 17 h after culture. Kinetochores were stained with CREST (red) and chromosomes with DAPI (blue). *Esrp1*-knockout oocytes had entered to metaphase II with univalent sister chromatids after reversine treatment. Insets show typical chromosome figures observed. Scale bar, 1 µm.

### Preferential nuclear-localized ESRP1 is involved in oocyte mRNA processing

We next used oocytes from PD5, as well as PD12 ovaries and adolescent mice pre-treated with PMSG for 44–46 h that were allowed to mature *in vitro* for 0, 8, and 14 h (corresponding to the GV-NSN/SN, MI, and MII stages, respectively), to examine the subcellular localization of ESRP1 during each oocyte maturation stage. ESRP1 was found to be preferentially localized to the nucleus in PD5, PD12, and GV-NSN oocytes when transcription was active, but relatively low ESRP1 levels were found in GV-SN, MI, and MII oocytes when transcription was inactive (Fig. 7A). Next, using alkyne-modified nucleoside 5-ethynyl uridine (5-EU), which reflects newly synthesized RNAs and global RNA synthesis, we found that 5-EU staining in NSN-type oocytes was colocalized with high levels of ESRP1 in the nucleus, whereas 5-EU was undetectable in SN-type oocytes, and low levels of ESRP1 were found in the cytoplasm (Fig. 7B), suggesting that ESRP1 was involved in mRNA processing during oocyte maturation.

**Figure 7.**
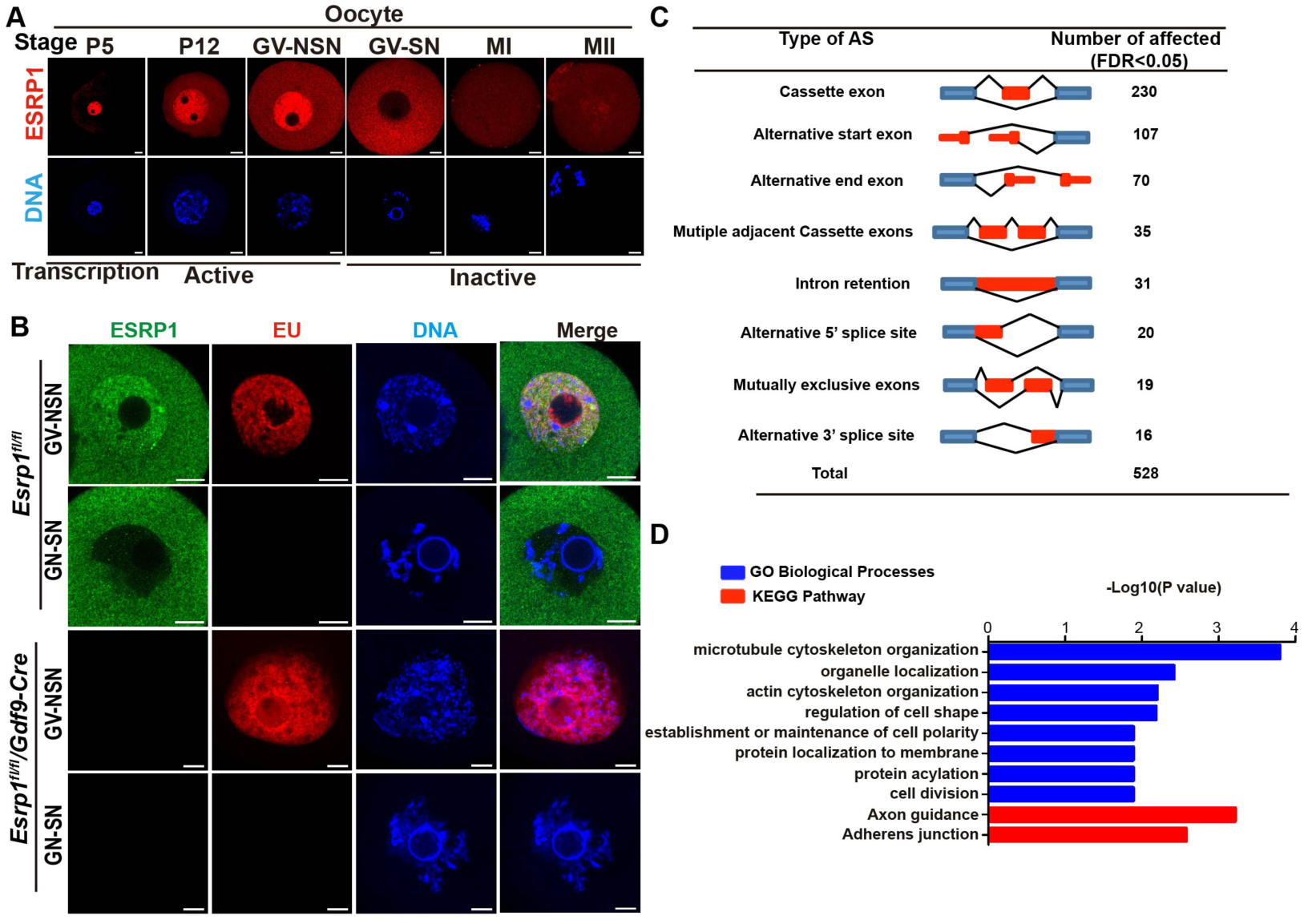
Preferential nuclear-localized ESRP1 is involved in oocyte mRNA processing. (A) Immunofluorescence showing localization of ESRP1 (red) in oocytes. DNA was stained with DAPI (blue). Abbreviations: P5, P12: postnatal days 5, 12; GV: germinal vesicle stage; NSN: non-surrounded nucleolus; SN: surrounded nucleolus; MI: metaphase I; MII, metaphase II. Scale bars, 10 µm. (B) Co-staining of 5-EU (red) and ESRP1 (green) in mouse oocytes in NSN- and SN-nucleolus types at GV stages. DNA is shown in blue. Scale bars, 10 µm. (C) Diagrams of eight AS events recognized by CASH software and splicing events were significantly affected by the depletion of *Esrp1* in ovulated oocytes analyzed via RNA-seq data using CASH software (FDR<0.05). (D) GO-term enrichment and KEGG-pathway analyses of genes with significantly affected AS events.

We next investigated the regulatory effects of ESRP1 and global RNA changes due to ESRP1 knockout in mouse oocytes. Transcriptomes were compared in ovulated oocytes between six-week-old *Esrp1*^*fl/fl*^ and *Esrp1*^*fl/fl*^*/Gdf9-Cre* females via high-throughput sequencing (RNA-seq; four biological replicates in each group, GEO database GSE149355). Approximately 85% of clean reads were applied for analyses (Fig. S4A), and proximal values of global gene expression were shown within samples (Fig. S4B). We found that 3,002 genes (1586 downregulated and 1416 upregulated) showed significant differences (FDR value < 0.01, more than 1.5-fold and baseMeand ⩾10) in transcript levels between *Esrp1*^*fl/fl*^ and *Esrp1*^*fl/fl*^*/Gdf9-Cre* (Table S3). Among them, numerous oocyte meiosis-related genes were downregulated in *Esrp1-*knockout ovulated oocytes including *Ccnb1 (cyclin B1), Ccnb3 (cyclin B3), Marf1, Pcnt, Dazl, Kif1b, Rab8b*, and *Spire1*, while some genes involved in anaphase-I transformation—such as *Bub3* and *Pttg1 (Securin)*—were upregulated (Fig. S4C). We further validated mRNA levels via quantitative real-time PCR (qRT-PCR) in oocytes (Fig. S4D). We found that mRNA levels of genes important for oocyte meiosis were significantly altered due to the conditional deletion of *Esrp1* in mouse oocytes.

ESRP1 has been reported to functionally regulate AS (Warzecha et al. 2009b; Bebee et al. 2015; Mager et al. 2017). Hence, we next analyzed global AS events between *Esrp1*^*fl/fl*^ and *Esrp1*^*fl/fl*^*/Gdf9-Cre* oocytes via comprehensive AS hunting method (CASH) (Wu et al. 2018). We revealed that 528 AS events from a total of 76,116 splicing events were affected in *Esrp1-*knockout oocytes (FDR value < 0.05, Table S4). Our analysis revealed that the major cluster of splicing events was cassette exon (230) among the 528 AS events, while others included alternative start exon (107), alternative end exon (70), multiple adjacent cassette exons (35), intron retention (31), alternative 5’ splice site (20), mutually exclusive exons (19), and alternative 3’ splice site (16) (Fig. 7C). These AS events occurred in a total of 459 genes and were subjected to Gene Ontology analysis. Microtubule cytoskeletal organization was the top-affected biological process following *Esrp1* deletion, along with organelle localization, actin cytoskeletal organization, and establishment/maintenance of cellular polarity (Fig. 7D). Similarly, Kyoto Encyclopedia of Genes and Genomes (KEGG) pathway analysis further suggested that abnormal spindle organization in *Esrp1-*knockout oocytes was likely due to aberrant AS in genes associated with cytoskeletal organization (Fig. 7D).

### Pre-mRNA splicing of spindle-organization-associated genes is required for oocyte meiosis

Using GV-stage and ovulated oocytes from *Esrp1*^*fl/fl*^ and *Esrp1*^*fl/fl*^*/Gdf9-Cre* females, we next looked for splicing events due to *Esrp1* deletion. We confirmed that the three exons of *Esrp1* were deleted completely in oocytes, as indicated in *Gdf9-Cre* mice (Fig. 8A). A previous study reported that ESRP1 mediates splicing of *Fgfr2* pre-mRNA (Warzecha et al. 2009a). In our present study, an identical splicing defect of *Fgfr2* (termed mutually exclusive exons) was found in *Esrp1*^*fl/fl*^*/Gdf9-Cre* oocytes (Fig. 8A). Microtubule cytoskeletal organization has been shown to represent the key step during mouse oocyte meiosis to maintain normal chromosome alignment and spindle formation (Bennabi et al. 2016), and it is the most affected cellular process due to deletion of *Esrp1* in oocytes. Therefore, we next selected nine genes that have previously been shown to be associated with spindle organization during oocyte meiosis for further analysis. We employed Integrative Genomics Viewer (IGV), a visualization tool for efficient and flexible exploration of large and complicated data sets obtained from sequencing (Robinson et al. 2011), to analyze ESRP1-mediated AS of these nine genes. Among these AS events, *Trp53bp1, Rac1, Bora, kif23, Ndel1, kif3a, Cenpa*, and *Lsm14b* were classified as cassette exon, whereas *kif2c* was an alternative 5’ splice site (Fig. 8A). Next, we used specific primers (Table S1) of target genes and semi-quantitative PCR to determine the aberrant splicing patterns of these nine genes in GV-stage and ovulated oocytes from *Esrp1*^*fl/fl*^ and *Esrp1*^*fl/fl*^*/Gdf9-Cre* females (Fig. 8B), which represent two key developmental stages during mouse oocyte development. Among the nine genes, the splicing changes of *Lsm14b* were the most obvious according to IGV and CASH analyses (Fig 8A and Table S4); LSM14B directly binds to tubulin, and is essential for spindle stability and oocyte meiosis (Mili et al. 2015; Zhang et al. 2017), so we use antibody to verify the protein change. Strikingly, as revealed by Western blotting, ESRP1 depletion led to an obvious shift from the standard isoform of LSM14B to the longer isoform of LSM14B (Fig. 8C). In *Esrp1*^*fl/fl*^ oocytes, there were two ESRP1-dependent splicing events in *Lsm14b*, including the sixth exon (predominant) and the fourth exon skipped (Fig. 8A, B); however, it was retained in *Esrp1*^*fl/fl*^*/Gdf9-Cre* oocytes (Fig. 8A, B). In *Esrp1*^*fl/fl*^ mice, both isoforms of LSM14B proteins were found in GV-stage oocytes, and the standard isoform was predominant. In contrast, in *Esrp1-*knockout oocytes, the protein levels of two isoforms of LSM14B were dramatically shifted at the GV stage. This shift was also confirmed in ovulated oocytes (Fig. 8C). The expression of the LSM14B standard isoform was dramatically decreased by 86%(Fig. 8D), whereas expression of the longer isoform was increased by 145% in GV-stage oocytes in *Esrp1-*knockout oocytes (Fig. 8E). Taken together, these data suggest that the global ESRP1-mediated AS program was required for maintaining spindle morphogenesis and oocyte meiosis.

**Figure 8.**
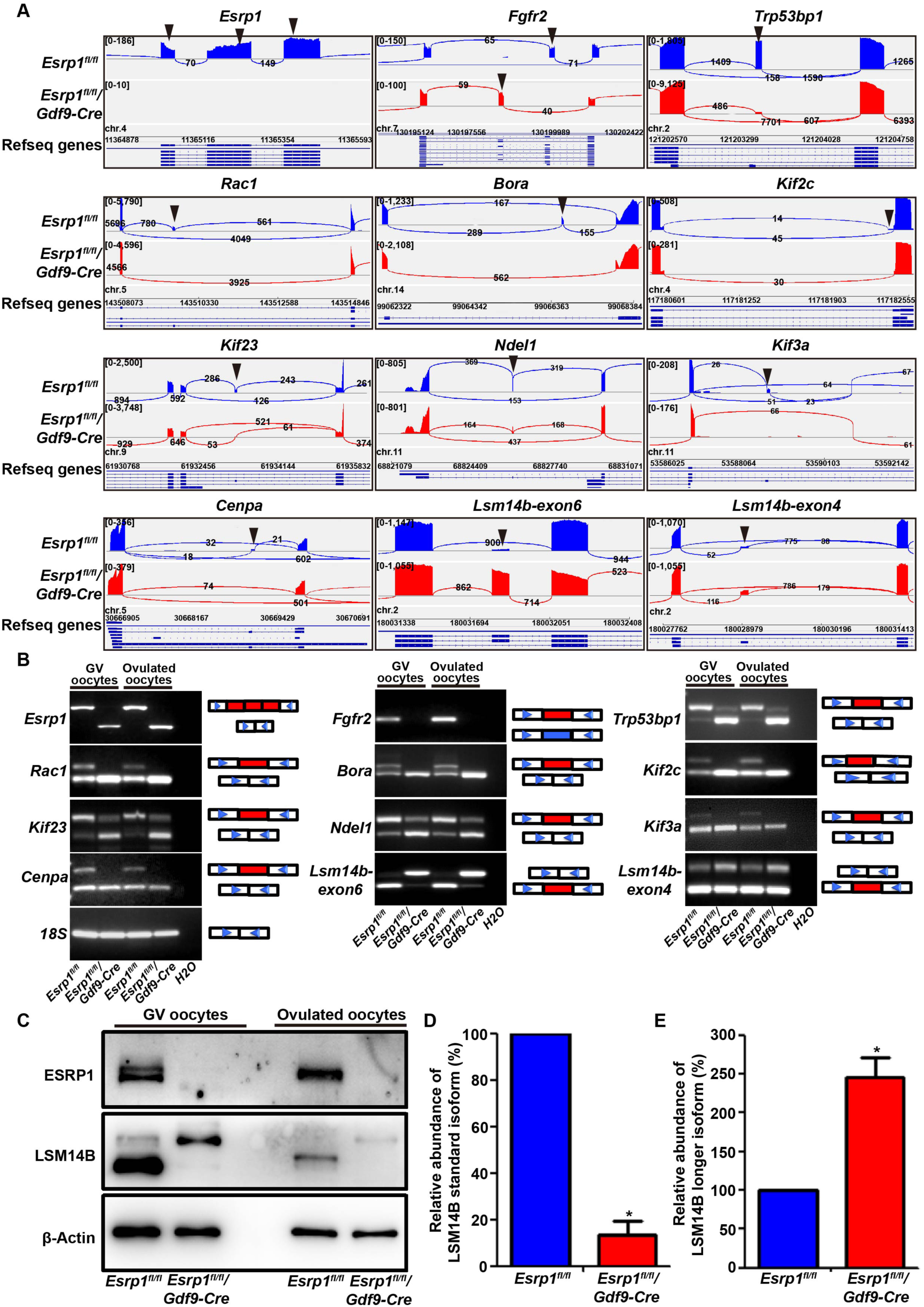
Loss of ESRP1 in oocytes leads to global changes in pre-mRNA splicing. (A) Use of IGV software shows alternative sites in genes related to oocyte spindle organization, as indicated from RNA-seq results. Black arrowheads indicate splicing sites. (B) PCR analysis of AS patterns of the changed spliced genes in GV-stage and ovulated stage oocytes from *Esrp1*^*fl/fl*^ and *Esrp1*^*fl/fl*^*/Gdf9-Cre* mice. 18s served as the internal control. PCR was performed with specific primers (Table S1) from three independent experiments. (C) Western-blot analyses showing the expression of two isoforms of LSM14B in GV-stage and ovulated stage oocytes from *Esrp1*^*fl/fl*^ and *Esrp1*^*fl/fl*^*/Gdf9-Cre* mice. β-actin served as a loading control. (D–E) Relative abundances of LSM14B standard and longer isoforms in GV-stage oocytes from *Esrp1*^*fl/fl*^ and *Esrp1*^*fl/fl*^*/Gdf9-Cre* mice were measured by Western-blot analyses from three independent experiments (*P < 0.05).

## Discussion

Our present study represents the first report to show that the splicing factor, ESRP1, controls oocyte meiosis and follicular development via pre-mRNA splicing in mice. We found that an abnormal splicing program resulting from ESRP1 deletion induced severe spindle organization defects, chromosome misalignment, abnormal SAC activation, and insufficient APC/C activity. These defects collectively resulted in first-meiosis arrest (Fig. 9). Since ESRP1 deletion hampered follicular development and ovulation, female infertility and ultimate POF were found in *Esrp1*-knockout mice.

**Figure 9.**
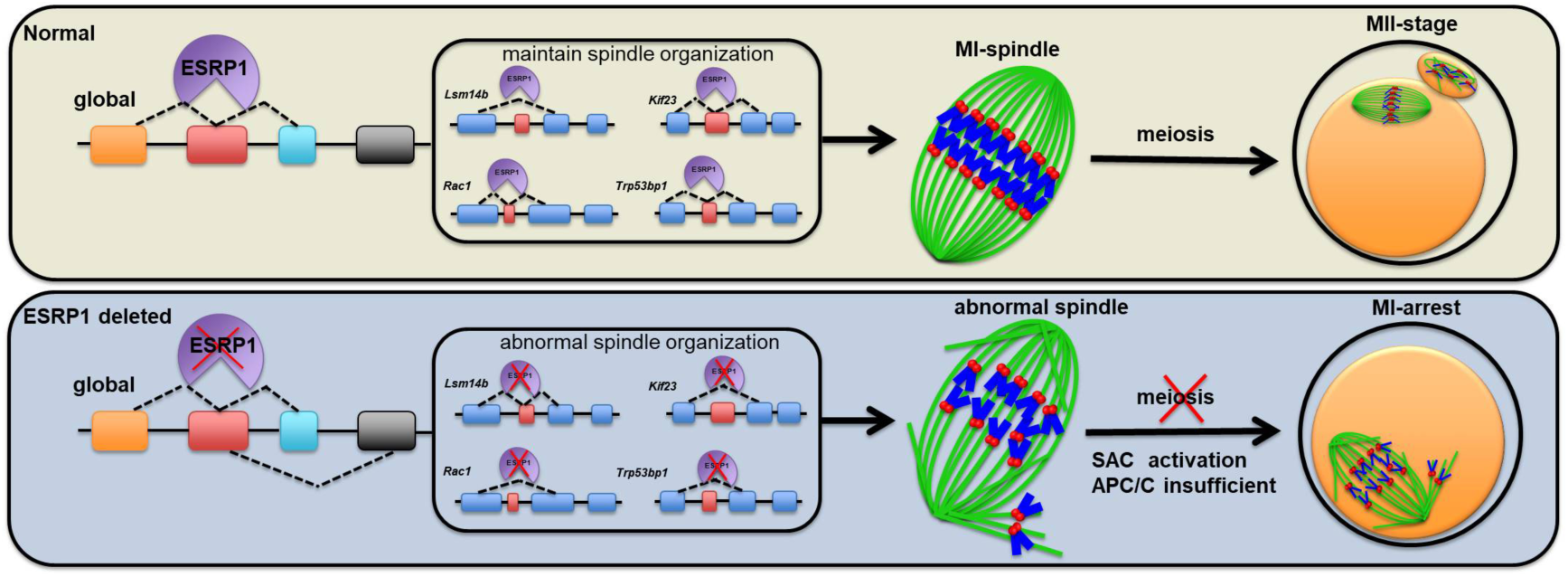
Proposed schematic model of ESRP1 function during oocyte meiotic progression. Generally, in normal oocytes, post-transcriptional modification of the maternal store mRNAs via AS and correct splicing promote to form typical barrel-shaped spindles and well-aligned chromosomes on the metaphase plate, leading to PBE and completion of the first meiosis. However, deletion of ESRP1 in oocytes leads to compromised global maternal mRNA splicing, which then leads to severe spindle organization defects, chromosomes misalignments, which contribute to abnormal activation of SAC and insufficiency of the APC/C. Hence, *Esrp1*-knockout oocytes failure to extrude the PB1 and are arrested at metaphase I.

Post-transcriptional modifications of mRNAs via AS are particularly critical in oocytes, as oocyte maturation is associated with suppression of transcription, and new mRNAs are not synthesized until after zygotic genome activation (ZGA) that occurs at the two-cell stage in mice (Flach et al. 1982). Gene transcription in the one-cell stage is uncoupled with splicing and 3’ processing, leaving most transcripts nonfunctional and genes transcribed in zygotes yield functional mRNAs until two-cell stage (Abe et al. 2015). During the period from oocyte maturation to ZGA, gene expression is mainly mediated by regulation of maternal mRNAs accumulated in the oocyte prior to meiotic reactivation (Christou-Kent et al. 2020; Sha et al. 2020).

ESRP1 is an RBP that regulates AS and has been found to be universally expressed in organisms that have been analyzed. Warzecha et al showed that ESRP1 binds to the *Fgfr2* auxiliary cis-element, intronic splicing enhancer/intronic splicing silencer-3 (ISE/ISS-3), to regulate splicing that is required for the expression of epithelial *Fgfr2*-IIIb. ISE/ISS-3 is located in the intron between mutually exclusive exons IIIb and IIIc, and ESRP1 promotes splicing of the upstream exon IIIb while silencing the downstream exon IIIc (Warzecha et al. 2009a). In our present study, mutually exclusive exons of *Fgfr2* were also found from our single-cell RNA-Seq data, consistent with the conventional action of ESRP1 in promoting exon inclusion by binding downstream of a regulated exon and possibly acting as an exon-skipping enhancer by binding to upstream introns (Warzecha et al. 2010). AS deficiency caused by ESRP1 has been reported in several mouse organs. ESRP1 enhances the expression of CD44v in breast cancer cells through AS, promoting breast cancer to lung metastasis and colonization independently of EMT (Yae et al. 2012); moreover, ablation of *Esrp1* results in ureteric branching defects and reduces nephron number due to incorrect *Fgfr2* splicing (Bebee et al. 2016). In the present study, ESRP1 was expressed in oocytes at all follicular stages. Since we found that ESRP1 was preferentially localized to the nucleus with active transcription at the NSN-GV stage, we investigated the function of ESRP1 during oocyte development and maturation in mice. Single-cell RNA-Seq analysis of *Esrp1*-knockout oocytes showed that ESRP1 determined the pre-mRNA processing of the oocyte maternal transcriptome. Thus, *Esrp1*^*fl/fl*^*/Gdf9-Cre* mice represent a model for investigating AS related to mouse meiosis. Previous work has shown that BCAS2 is involved in pre-mRNA splicing in spermatogonia in mouse testes. Disruption of *Bcas2* leads to a significant shift of isoforms of DAZL protein and impairs spermatogenesis and male mouse fertility (Liu et al. 2017). However, few studies have investigated the role of AS in female germ-cell development. Here, we revealed that ESRP1 mediated critical roles of the pre-mRNA splicing program necessary for female mouse fertility.

In the present study, ESRP1 deletion led to oocyte arrest at metaphase I, with abnormal spindle organization. In mouse oocytes, homologous chromosome separation requires separase to cleave the meiotic kleisin, REC8, and this step depends on APC/C-mediated degradation of cyclin B1 and securin (Herbert et al. 2003; Kudo et al. 2006). Furthermore, timely APC/C activation depends on satisfaction of the SAC (Holt et al. 2013; Touati and Wassmann 2016). In the present study, in *Esrp1*-knockout oocytes, defects in spindle organization and the presence of unattached chromosomes triggered SAC activity (Touati et al. 2015) and may have led to insufficient APC/C activity. Inhibition of SAC activity could partially rescue metaphase I arrest caused by *Esrp1* deficiency. Both activated SAC and insufficient APC/C activity may lead to metaphase-I arrest and affect the expression of numerous oocyte meiosis-related genes such as *Marf1* (Su et al. 2012).

Nevertheless, our present study showed that ESRP1 played pivotal roles in spindle/chromosome organization in mouse oocytes through AS. Maternal *Esrp1* depletion affects splicing of several hundred genes. Following these alterations in splicing, microtubule cytoskeletal organization was the most significantly affected process, and many ESRP1-controlled genes were associated with spindle organization. Mitotic kinesin-like protein 1 (MKlp1, also known as *Kif23*) contributes to the shortening and widening of the spindle during metaphase-I arrest in drosophila oocytes, and a partial depletion of MKlp1 produces significantly longer and narrower spindles in oocytes (Costa and Ohkura 2019). Tumor suppressor p53 binding protein 1 (*Trp53bp1*) plays a role in maintaining the bipolar spindle during mitosis (Lambrus et al. 2016). Recent studies have revealed that *Trp53bp1* knockdown affects spindle bipolarity and chromatin alignment by altering MTOC stability during oocyte maturation (Jin et al. 2019). In our present study, we also found abnormal splicing of *Rac1*, which has previously been shown to contribute to spindle positioning and oocyte cortical polarity during mouse oocyte meiosis (Halet and Carroll 2007). CENP-A, a variant of histone H3 specific for nucleosomes at centromeres, is required for the localization of all known kinetochore components. Preventing CENP-A deposition during prophase I-arrested oocytes results in a failure of kinetochore function, perturbs chromosome segregation, and ultimately cell dysfunction(Swartz et al. 2019). Mitotic centromere associated kinesin (MCAK; also known as *Kif2c*) is an important regulator of microtubule dynamics in a variety of systems, and knockdown of MCAK causes a delay in chromosome congression and induces a meiosis I arrest in mouse oocytes (Illingworth et al. 2010; Vogt et al. 2010). BORA is a critical regulator of Aurora A and Plk1, and a partial depletion of *Bora* affects mouse oocyte maturation and caused MI-arrest with spindle/chromosome disorganization and the disappearance of Aurora A and Plk1 at the spindle in oocytes(Zhai et al. 2013). Neurodevelopment protein 1-like 1 (*Ndel1*, also known as *NudEL*), is a dynein accessory protein. *Ndel1* mutant impairs the dynein-mediated transport of kinetochore proteins to spindle poles along microtubules, a process contributing to inactivation of the SAC in mitosis (Yan et al. 2003). KIF3A is involved in chromosomal structure maintenance (Monzo et al. 2012). Thus, a comprehensive set of abnormalities in splicing patterns of important genes regulated by ESRP1 may contribute to mouse meiotic arrest and infertility (Fig. 9).

LSM14 belongs to the RNA-associated protein (RAP) family and has been shown to exert roles in both mRNA storage and posttranscriptional regulation. Recent studies have shown that knockdown of LSM14B induces oocyte metaphase-I arrest and misalignment of chromosomes due to abnormal activation of SAC and MPF (Zhang et al. 2017). In the present study, from our single-cell sequencing data, we observed that the sixth exon of *Lsm14b* was spliced in *Esrp1*^*fl/fl*^ oocytes but was retained in *Esrp1*-knockout oocytes. Considering the profound effect of *Esrp1* loss on oocyte meiosis, the corresponding dramatic changes in *Lsm14b* have attracted particular interest. Here, we found that the protein level of the standard isoform of LSM14B was dramatically reduced, while the longer isoform of LSM14B was significantly up-regulated in GV-stage and ovulated oocytes from *Esrp1*^*fl/fl*^ */Gdf9-Cre* mice (Fig. 8C, D, E), suggesting that the standard isoform of LSM14B might represent the main functional form during mouse meiosis. Deletion of ESRP1 leads to dramatic retention of the sixth exon in LSM14B, suggesting a critical role for the precise control of *Lsm14b* splicing during normal mouse meiosis. However, the normal function of the longer isoform of LSM14B, with retention of the sixth exon, remains unknown. The dramatic decrease in LSM14B standard isoform and total LSM14B protein that we found in the present study suggests that deletion of ESRP1 leads to a significant degree of loss-of-function of LSM14B. In addition, a previous study has shown that LSM14B regulates the oocyte meiotic maturational process by controlling mRNA expression and degradation, such as of *Cyclin B1* and *Cdc20*, which are essential for the cell cycle (Zhang et al. 2017). Consistently, in our present study, we also confirmed that the mRNA levels of *Cyclin B1* and *Cdc20* were downregulated in GV-stage and ovulated oocytes from *Esrp1*^*fl/fl*^*/Gdf9-Cre* mice. These data suggest that ESRP1 might be critical for mouse meiosis via regulating *Lsm14b* splicing (Fig. 9). Collectively, our present findings seem to mimic the process in which AS defects cause abnormal oocyte meiosis, follicular development and ovulation disorders, and ultimate POF.

In summary, our present study is the first to elucidate that the RBP, ESRP1, is required for spindle organization, chromosome segregation during oocyte meiosis, and follicular development. Our findings shed light on the importance of AS in regulating female germline development and fertility and may also provide novel insights toward the clinical prediction of POF.

## Materials and Methods

### Animals

Mouse colonies in the present study were housed and maintained under specific-pathogen-free conditions in the Animal Core Facility of Nanjing Medical University (China). Mice were housed under a 12/12-h light/dark cycle, and food and water were provided *ad libitum*. All procedures and protocols involving mice were approved by the Institutional Animal Care and Use Committee (IACUC) at Nanjing Medical University (ID: IACUC1812016).

### Generation of *Esrp1*^*fl/Δ*^*/Ddx4-Cre and Esrp1*^*fl/fl*^ */Gdf9-Cre* mice

The targeting construct contained three *loxP* sites inserted into intron 6 and intron 9 of the *Esrp1* gene and a floxed HyTK double-selection cassette in exons 7–9. All three *loxP* sites were in the same orientation (Fig. S2A). The *Ddx4-Cre* (*Vasa-Cre*) mouse line was maintained on a mixed genetic background (129/C57BL/6×FVB/N) (cat. #J006954, Jackson Laboratory, Bar Harbor, ME). The Tg(Gdf9-iCre)5092Coo mouse line was maintained on the C57BL/6J genetic background (cat. #J011062, Jackson Laboratory). *Esrp1*^*fl/fl*^ mice were crossed with *Ddx4-Cre/+* or *Gdf9-Cre/+* transgenic mice to generate PGCs-specific *Esrp1*-knockout (*Esrp1*^*fl/Δ*^*/Ddx4-Cre)* mice or oocyte-specific *Esrp1*-knockout (*Esrp1*^*fl/fl*^ */Gdf9-Cre*) mice, respectively. Offspring was genotyped by PCR using specific primers (Table S1).

### Fertility analysis

For fertility assessment of *Esrp1*^*fl/Δ*^*/Ddx4-Cre* mice, 8-week-old *Esrp1*^*fl/fl*^ (control; n =6) and *Esrp1*^*fl/Δ*^*/Ddx4-Cre* mice (n = 6) were separately mated with WT C57BL/6 mice respectively at a ratio of 1:1. For *Esrp1*^*fl/fl*^ */Gdf9-Cre* mice, 8-week-old females (n = 6) were separately mated with WT C57BL/6 males at a ratio of 1:1. Mating pairs were continuously housed together for a period of six months, parturition frequency and litter sizes were assessed.

### Superovulation, oocyte collection, fertilization, and embryo culture

Most mice, unless specify, used in this study were 4-6-week-old. For the superovulation assay, mice were intraperitoneally injected with 5 international units (IU) of PMSG (lot. #190915, Ningbo Sansheng Pharmaceutical, CHN) to stimulate antral follicular development and 44**–** 46 h later with another 5 IU of hCG (lot. #180428, Ningbo Sansheng Pharmaceutical, CHN), 14**–**16 h later, cumulus-oocyte complexes (COCs) were surgically disassociated from oviducts and digested with 0.5mg/ml hyaluronidase (cat. #H4272, Sigma-Aldrich, USA), and the total ovulated oocyte number was counted. For oocyte collection, mice were prepared by intraperitoneal injection of 5 IU of PMSG 44**–** 46 h before oocyte collection. Fully-grown oocytes arrested at germinal vesicle (GV) stage were collected from ovaries in M2 medium (Cat. #M7167, Sigma-Aldrich) with 5 µM Milrinone (Cat. #475840, Millipore, USA). Cumulus cells were removed by gently pipetting. For IVM, oocytes with the intact GV were selected for culture in the M2 medium under paraffin oil at 37°C in a 5% CO_2_ atmosphere. Cells were removed from culture at the appropriate stages or time points for the following assays. The progress of meiosis in the oocytes was determined based on the ratio of GVBD and PBE. For Mps1 inhibition, reversine (cat. #10004412, Cayman Chemical Research) was added to the culture medium 5.5 h after GVBD at a final concentration of 0.5 µM. As oocytes are cultured in drops covered with mineral oil and because reversine is soluble in oil, mineral oil covering the culture drops contained reversine as well, and untreated controls were cultured in separate dishes. For IVF, normal sperm isolated from B6D2F1 adult males were inseminated with the ovulated oocytes with visible PB1 or COCs. Formation of 2-cell stage embryos was scored 24 h after IVF, and 2-cell stage embryos were transferred into KSOM (cat. #MR-020P-5F, Millipore, USA) medium for further development. Development of 2-cell embryos to the blastocyst stage was scored 3**–**5 days after culture. Oocytes and embryos morphology were observed and images were acquired with a stereoscope (1×51/0N3-M300, Olympus, Germany).

### Histology, immunofluorescence, and immunohistochemistry

Tissues were fixed in Modified Davidson’s Fluid (MDF) overnight at room temperature and were then embedded in either paraffin—for histological and immunohistochemical analyses—or in 4% paraformaldehyde (PFA, cat. #P6148, Sigma-Aldrich) overnight at 4°C, after which tissues were embedded in optimal-cutting-temperature compound (OCT, cat. #4583, Sakura) for immunofluorescence. Paraffin sections (5 µm) were stained with hematoxylin and eosin (HE) and observed under a microscope (Axio Scope A1, Carl Zeiss, Germany). Follicles at different developmental stages were counted in every third section throughout the whole ovary. To avoid repeated counting of the same follicle, only follicles containing an oocyte with visible GV were counted. The stages of follicles were categorized as follows: primordial follicles were surrounded by a layer of partially or intact squamous granules; primary follicles were surrounded by a layer of cuboidal granulosa cells; early-secondary follicles had less than or equal to two layers of granulosa cells; later-secondary follicles had more than two layers of granulosa cells without visible antrum; and antral follicles had fluid-filled antrum forms. For immunofluorescence and immunohistochemistry, following hydration and antigen retrieval, tissue sections (5 µm) were permeated in PBS containing 0.3% Triton X-100 for 15 min, blocked with 3% bovine serum albumin (BSA, cat. #A3803,

Sigma-Aldrich) in PBS for 2 h at room temperature, and were incubated with primary antibodies (Table S2) diluted in 3% BSA overnight at 4°C. After washing three times with PBS, sections were incubated with TRITC/FITC-conjugated secondary antibody (Table S2) for immunofluorescence or with horseradish peroxidase (HRP)-conjugated secondary antibody (Table S2) and diaminobenzidine (DAB, cat. #ZLI9017, Zhongshan Jinqiao biotechnology) for immunohistochemistry. The sections were then washed with PBS three times, incubated with DAPI (1µg/ml in PBS, cat. #D9542, Sigma-Aldrich) for 10 min at room temperature, and observed using a confocal laser-scanning microscope (LSM 800, Carl Zeiss, Germany) or ZEISS Axio Scope A1. For oocytes immunofluorescence, oocytes were fixed in 4% PFA for 30 min at room temperature, permeated in PBS containing 0.5% Triton X-100 for 20 min at room temperature, blocked with 1% BSA in PBS for 1 h at room temperature and were incubated with the primary antibody (Table S2) overnight at 4°C. Following washing three times with PBST (0.1% Tween 20 and 0.01% Triton X-100 in PBS), oocytes were incubated with appropriate secondary antibody (Table S2) for 1 h at room temperature. After washing three times, oocytes were stained with DAPI (1µg/ml in PBS) for 10 min. For K-MT attachments assessment, oocytes were briefly chilled at 4°C to induce depolymerization of non-kinetochore microtubules just prior to fixation. Finally, oocytes were mounted on slides in an anti-fade reagent (cat. #P0126, Beyotime, CHN), and observed using a confocal laser-scanning microscope (LSM 800, Carl Zeiss, Germany).

### 5-Ethynyl uridine (EU) staining

The 5-ethynyl uridine (5-EU) staining was performed with a Cell-Light EU Apollo567 RNA Imaging Kit (cat. #C00031, RiboBio, CHN) according to the manufacturer’s instructions, with the exception that 5-EU was added to the M2 medium with 5 µM of milrinone and was incubated with GV oocytes for 2 h. Subsequently, immunofluorescence was performed as described above.

### Single-cell RNA sequencing

A total of eight oocyte RNA samples were used from six-week-old *Esrp1*^*fl/fl*^ and *Esrp1*^*fl/fl*^*/Gdf9-Cre* female mice according to four individual collections. One in every 15 oocytes was collected in a tube with ribonuclease inhibitor (cat. #Y9240L, Enzymatics, GER) and lysis buffer. The directly lysed oocytes were used for cDNA synthesis using Smart-seq2. After amplification, the cDNA was fragmented by ultrasonic waves using a Bioruptor Sonication System (Diagenode Inc., Denville, USA). Sequencing libraries were made with the fragmented DNA using a NEBNext Ultra DNA Library Prep Kit for Illumina according to the manufacturer’s instructions (cat. #E7370, New England Biolabs, USA). Library quality inspection was established via a LabChip GX Touch (PerkinElmer) and Step OnePlus Real-Time PCR System (ThermoFisher, USA). Qualified libraries were then loaded on a HiSeq X Ten platform (Illumina, San Diego, USA) for paired-end 150-bp sequencing.

### RNA extraction and reverse-transcription PCR (RT-PCR)

Total RNA was extracted using RNeasy Micro Kits (cat. #74004, Qiagen, GER) according to the manufacturer’s instructions. Reverse transcription (RT) was performed using the PrimeScript RT master mix (cat. #RR036A, Takara, Japan). Real-time RT-PCR was performed in a StepOne plus system (ABI StepOne Plus, ThermoFisher, USA) using SYBR green master mix (cat. #Q141-02, Vazyme, China). Threshold cycle (*Ct*) values were obtained, and the *ΔΔCt* method was used to calculate the fold changes. All of the values were normalized to 18s. Primer sequences are listed in Table S1.

### Protein extraction and western blot

Oocytes were collected from *Esrp1*^*fl/fl*^ and *Esrp1*^*fl/fl*^ */Gdf9-Cre* female at indicated time points, and washed in PBS-PVA to remove proteins from the medium. All samples were lysed in RIPA buffer (cat. #P0013C, Beyotime, CHN) containing protease inhibitor cocktail (cat. #11697498001, Roche, USA). The samples were loaded into 5×SDS loading buffer (cat. #P0015L, Beyotime, CHN) and boiled for 5 min. The lysates were subjected to 5% Sodium dodecyl sulfate polyacrylamide gel electrophoresis (SDS-PAGE) stacking gel at 80 V and a 10% SDS-PAGE separating gel at 120 V. The separated proteins were transferred onto polyvinylidene fluoride (PVDF) membranes (cat. #162-0177, Bio-Rad, USA). Thereafter, membranes were blocked in 5% skimmed milk in TBST (Tris buffered saline containing 0.1% Tween-20) for 2 h at room temperature and incubated with primary antibody (Table S2) overnight at 4°C. After washing with TBST three times, the membranes were incubated with a secondary antibody conjugated with HRP (Table S2) for 2h at room temperature, followed by stringent washing. Protein bands were detected by using ECL reagent (NEL105001EA, Perkinelmer, USA).

### Chromosome spreads

For chromosome spreads, oocytes were harvested from *Esrp1*^*fl/fl*^ and *Esrp1*^*fl/fl*^ */Gdf9-Cre* female. Oocytes were exposed to acid Tyrode’s solution (cat. #T1788, Sigma-Aldrich, USA) for 1-2 min at 37 °C to remove zona pellucidae. Next, the oocytes were recovered in M2 medium for 10 min, and fixed in a drop of spread solution with 1% PFA, 0.15% Triton X-100, and 3mM DTT (cat. #R0861, Thermo Scientific, USA) on the clean glass slides. After air-drying, the slides were blocked with 1% BSA in PBS for 1h at room temperature and incubated with the primary antibody (Table S2) overnight at 4°C, then incubated with corresponding secondary antibodies (Table S2) for 1 h at at room temperature. DNA counterstaining were stained with DAPI (1µg/ml in PBS) and observed using a confocal laser-scanning microscope (LSM 800, Carl Zeiss, Germany).

### Statistical analysis

All experiments were performed at least three times independently. All quantitative data are presented as the mean ± the standard error of mean (SEM), and analyses were carried out via GraphPad Prism 8.00 (GraphPad Software, Inc.). Statistical differences between each group were analyzed by two-tailed Student’s *t* tests or rank-sum tests via SPSS software 19.0 (IBM Corporation). A *P* value < 0.05 was considered to be statistically significant.

### Data availability

RNA sequencing data reported in this paper are accessible through the NCBI Gene Expression Omnibus (GEO) accession number GSE149355.

## Acknowledgments

We would like to thank CAM-SU International Cooperation Center, Soochow University, China for help in generation of *Esrp1-floxed* mice. We are grateful to all female fertility research groups (Dr YQ Su, J Li, Q Wang and D Zhang) at SKLRM for their valuable comments and technical help. We thank to Dr Ralph L Brinster for his support on the finding of RBP profiling in germline cells (Dr Xin Wu was a previous postdoc from Brinster lab) and Dr Jeremy P Wang at University of Pennsylvania for the suggestion and support for the RBP studies (Dr Jian Zhou is a previous postdoc from Wang lab). We thanks to LetPub for help with manuscript grammar checking. We also thank all valuable helps from the members in Wu laboratory. The work was supported by the National Key R&D Program of China (2018YFC1003302 to XW), the National Natural Science Foundation of China (31872844 to XW), the National Natural Science Foundation of Jiangsu Province (BK20181134 to JZ).

## Author contributions

X.W and J.Z conceived the study. L.P.Y., X.B.G., and H.R.Z. bred the mice. L.P.Y., and H.R.Z. performed most of the experiments, J.Z. constructed the ES cell targeting. L.P.Y., and D.D.Q. designed the primers to verify the aberrant splicing patterns. L.P.Y. prepared the figures. L.P.Y. and X.W wrote the manuscript. The manuscript was reviewed by all authors.

## Figures and legends

**Figure S1.**
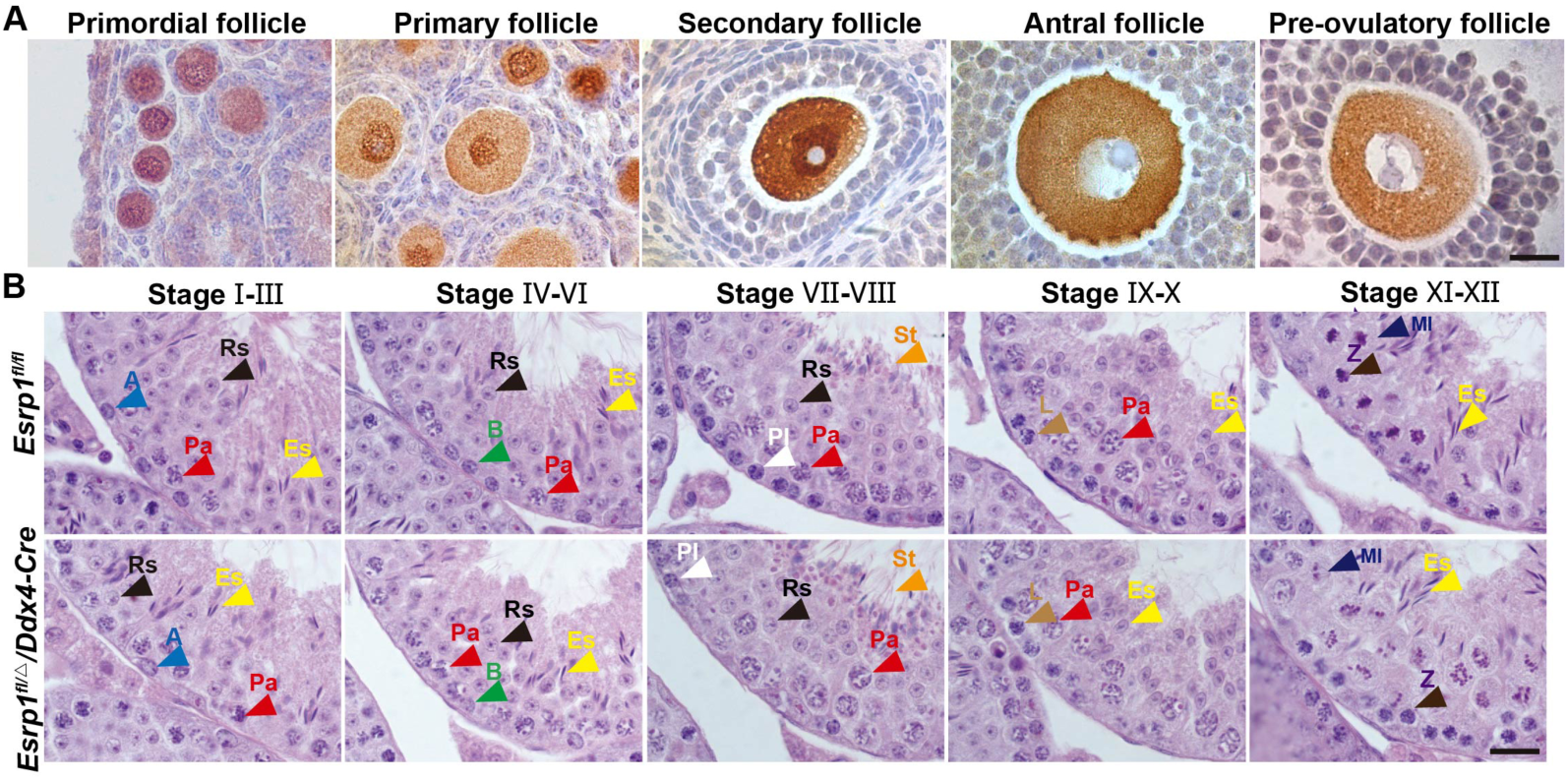
Expression and subcellular localization of ESRP1 during follicular development. (A) An immunohistochemistry (IHC) of ESRP1 in ovarian sections showing ESRP1 expression in follicles at indicated developmental stages. Scale bars, 20 µm. (B) HE staining of adult *Esrp1*^*fl/fl*^ and *Esrp1*^*fl/Δ*^*/Ddx4-Cre* testicular sections. Scale bars, 20 µm. Abbreviations: A: Type A spermatogonia; B: Type B spermatogonia; PI: Preleptotene spermatocyte; L: Leptotene spermatocyte; Z: Zygotene spermatocyte; Pa: Pachytene spermatocyte; MI: Post-prophase I meiotic phase; Rs: Round spermatid; Es: Elongated spermatid; St: spermatozoa.

**Figure S2.**
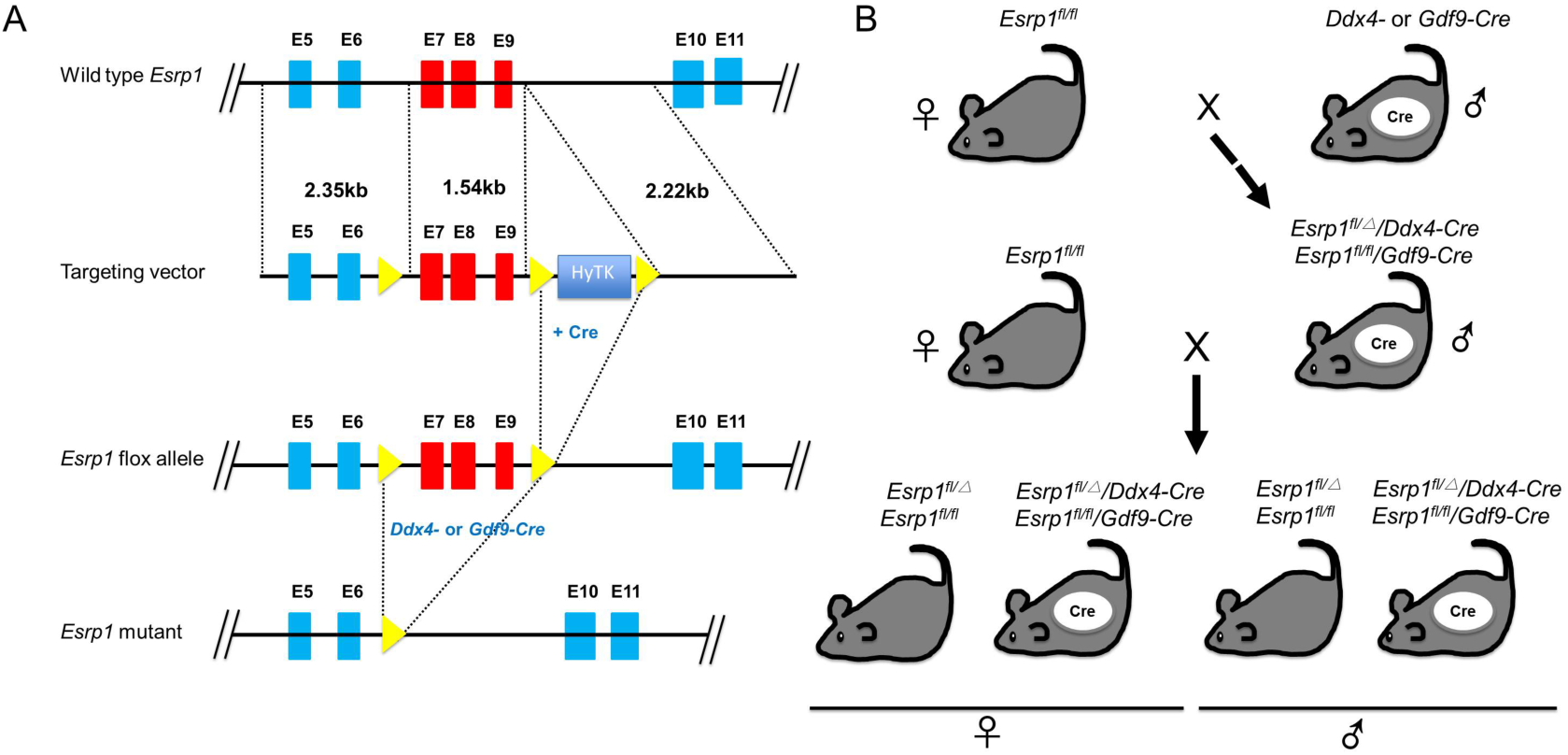
Generation of *Esrp1* conditional-knockout mice. (A) Construction of *Esrp1*^*Flox/Flox*^ and generation of *Esrp1*^*fl/Δ*^*/Ddx4-Cre* and *Esrp1*^*fl/fl*^ */Gdf9-Cre* mice. Exons 7–9 of *Esrp1* were deleted by *Ddx4-Cre*-mediated or *Gdf9-Cre*-mediated recombination. (B) Schematic of the breeding scheme to produce PGCs-specific *Esrp1* knockout in *Esrp1*^*fl/Δ*^*/Ddx4-Cre* mice and oocyte-specific *Esrp1* knockout in *Esrp1*^*fl/fl*^ */Gdf9-Cre* mice.

**Figure S3.**
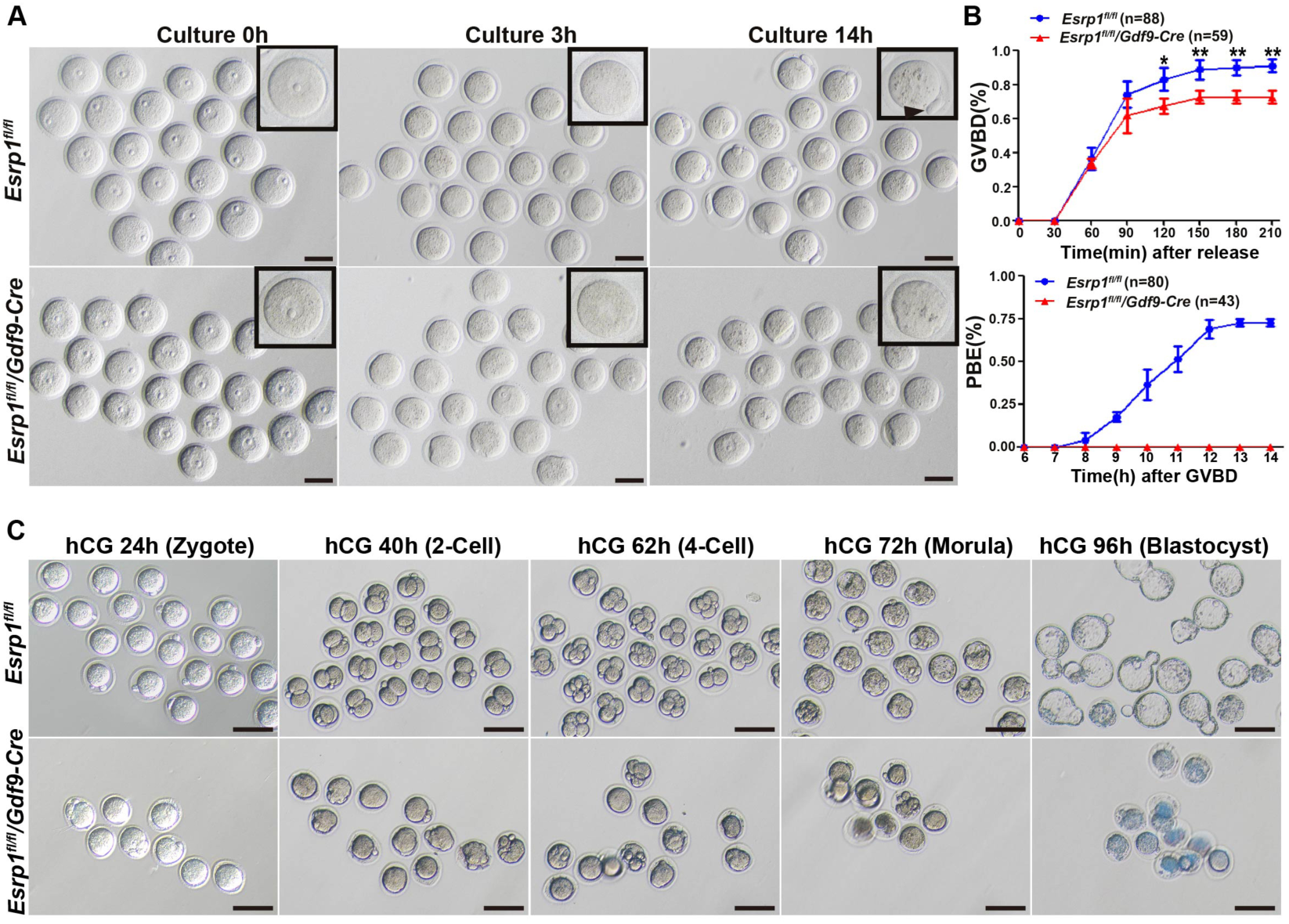
Compromised meiosis and embryonic development in *Esrp1*-knockout oocytes. (A) DIC imaging of the progression of meiosis in *Esrp1*^*fl/fl*^ and *Esrp1*^*fl/fl*^*/Gdf9-Cre* oocytes in culture. Time indicates hours after oocytes were released from meiotic arrest. Scale bar, 100 mm. (B) GVBD and PBE rates of cultured oocytes shown in (A). n: number of used oocytes. (C) DIC imaging of *Esrp1*^*fl/fl*^ and *Esrp1*^*fl/fl*^*/Gdf9-Cre* female embryos cultured *in vitro*. Embryonic development was monitored at the indicated time points after hCG administration. Scale bar, 100 µm.

**Figure S4.**
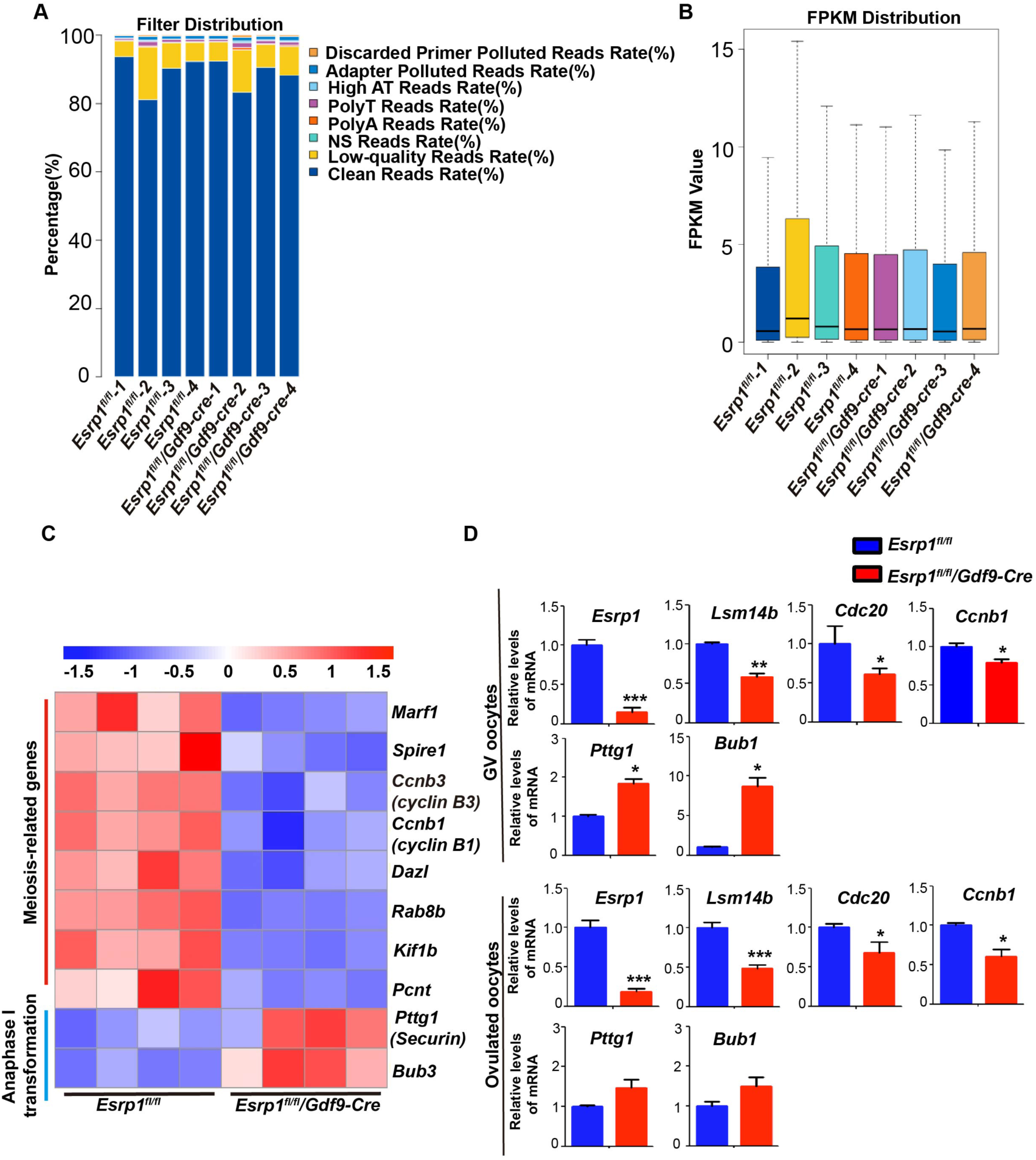
ESRP1 regulates mRNA transcription involved in oocytes meiosis. (A) Filter distribution in each sample; the clean reads were filtered for analyses. (B) Distribution of global gene expression in each sample. (C) A heatmap showing the transcript level of key genes that were involved in meiosis in *Esrp1*^*fl/fl*^ and *Esrp1*^*fl/fl*^*/Gdf9-Cre* oocytes. The color indicates the expression level. (D) Real-time RT-PCR analysis of mRNA levels of a cohort of genes. The mRNA levels of *Esrp1, Lsm14b, cdc20*, and *Ccnb1* were decreased in both GV-stage and ovulated oocytes from *Esrp1*^*fl/fl*^*/Gdf9-Cre* mice compared to those in *Esrp1*^*fl/fl*^ mice. The mRNA levels of *Bub1*and *Pttg1* were increased in both GV-stage and ovulated oocytes from *Esrp1*^*fl/fl*^*/Gdf9-Cre* mice compared to those in *Esrp1*^*fl/fl*^ mice. (\**p* < 0.05, \*\**p* < 0.01 and\*\*\**p* < 0.001).

## Reference

Abe K, Yamamoto R, Franke V, Cao M, Suzuki Y, Suzuki MG, Vlahovicek K, Svoboda P, Schultz RM, Aoki F. 2015. The first murine zygotic transcription is promiscuous and uncoupled from splicing and 3’ processing. The EMBO journal 34: 1523–1537.

Bebee TW, Park JW, Sheridan KI, Warzecha CC, Cieply BW, Rohacek AM, Xing Y, Carstens RP. 2015. The splicing regulators Esrp1 and Esrp2 direct an epithelial splicing program essential for mammalian development. eLife 4.

Bebee TW, Sims-Lucas S, Park JW, Bushnell D, Cieply B, Xing Y, Bates CM, Carstens RP. 2016. Ablation of the epithelial-specific splicing factor Esrp1 results in ureteric branching defects and reduced nephron number. Developmental dynamics : an official publication of the American Association of Anatomists 245: 991–1000.

Bennabi I, Terret ME, Verlhac MH. 2016. Meiotic spindle assembly and chromosome segregation in oocytes. J Cell Biol 215: 611–619.

Black DL. 2000. Protein diversity from alternative splicing: a challenge for bioinformatics and post-genome biology. Cell 103: 367–370.

Brown RL, Reinke LM, Damerow MS, Perez D, Chodosh LA, Yang J, Cheng C. 2011. CD44 splice isoform switching in human and mouse epithelium is essential for epithelial-mesenchymal transition and breast cancer progression. The Journal of clinical investigation 121: 1064–1074.

Christou-Kent M, Dhellemmes M, Lambert E, Ray PF, Arnoult C. 2020. Diversity of RNA-Binding Proteins Modulating Post-Transcriptional Regulation of Protein Expression in the Maturing Mammalian Oocyte. Cells 9.

Costa MFA, Ohkura H. 2019. The molecular architecture of the meiotic spindle is remodeled during metaphase arrest in oocytes. J Cell Biol 218: 2854–2864.

Do DV, Strauss B, Cukuroglu E, Macaulay I, Wee KB, Hu TX, Igor RLM, Lee C, Harrison A, Butler R et al. 2018. SRSF3 maintains transcriptome integrity in oocytes by regulation of alternative splicing and transposable elements. Cell discovery 4: 33.

Du G, Wang X, Luo M, Xu W, Zhou T, Wang M, Yu L, Li L, Cai L, Wang PJ et al. 2020. mRBPome capture identifies the RNA-binding protein TRIM71, an essential regulator of spermatogonial differentiation. Development (Cambridge, England) 147.

Flach G, Johnson MH, Braude PR, Taylor RA, Bolton VN. 1982. The transition from maternal to embryonic control in the 2-cell mouse embryo. The EMBO journal 1: 681–686.

Graveley BR, Brooks AN, Carlson JW, Duff MO, Landolin JM, Yang L, Artieri CG, van Baren MJ, Boley N, Booth BW et al. 2011. The developmental transcriptome of Drosophila melanogaster. Nature 471: 473–479.

Halet G, Carroll J. 2007. Rac activity is polarized and regulates meiotic spindle stability and anchoring in mammalian oocytes. Dev Cell 12: 309–317.

Hentze MW, Castello A, Schwarzl T, Preiss T. 2018. A brave new world of RNA-binding proteins. Nat Rev Mol Cell Biol 19: 327–341.

Herbert M, Levasseur M, Homer H, Yallop K, Murdoch A, McDougall A. 2003. Homologue disjunction in mouse oocytes requires proteolysis of securin and cyclin B1. Nat Cell Biol 5: 1023–1025.

Hodges CA, Hunt PA. 2002. Simultaneous analysis of chromosomes and chromosome-associated proteins in mammalian oocytes and embryos. Chromosoma 111: 165–169.

Holt JE, Lane SI, Jones KT. 2013. The control of meiotic maturation in mammalian oocytes. Current topics in developmental biology 102: 207–226.

Illingworth C, Pirmadjid N, Serhal P, Howe K, Fitzharris G. 2010. MCAK regulates chromosome alignment but is not necessary for preventing aneuploidy in mouse oocyte meiosis I. Development (Cambridge, England) 137: 2133–2138.

Jin ZL, Suk N, Kim NH. 2019. TP53BP1 regulates chromosome alignment and spindle bipolarity in mouse oocytes. Molecular reproduction and development 86: 1126–1137.

Kageyama S, Liu H, Kaneko N, Ooga M, Nagata M, Aoki F. 2007. Alterations in epigenetic modifications during oocyte growth in mice. Reproduction 133: 85–94.

Kasowitz SD, Ma J, Anderson SJ, Leu NA, Xu Y, Gregory BD, Schultz RM, Wang PJ. 2018. Nuclear m6A reader YTHDC1 regulates alternative polyadenylation and splicing during mouse oocyte development. PLoS genetics 14: e1007412.

Kudo NR, Wassmann K, Anger M, Schuh M, Wirth KG, Xu H, Helmhart W, Kudo H, McKay M, Maro B et al. 2006. Resolution of chiasmata in oocytes requires separase-mediated proteolysis. Cell 126: 135–146.

Lambrus BG, Daggubati V, Uetake Y, Scott PM, Clutario KM, Sluder G, Holland AJ. 2016. A USP28-53BP1-p53-p21 signaling axis arrests growth after centrosome loss or prolonged mitosis. J Cell Biol 214: 143–153.

Lan ZJ, Xu X, Cooney AJ. 2004. Differential oocyte-specific expression of Cre recombinase activity in GDF-9-iCre, Zp3cre, and Msx2Cre transgenic mice. Biol Reprod 71: 1469–1474.

Liu W, Wang F, Xu Q, Shi J, Zhang X, Lu X, Zhao ZA, Gao Z, Ma H, Duan E et al. 2017. BCAS2 is involved in alternative mRNA splicing in spermatogonia and the transition to meiosis. Nat Commun 8: 14182.

Mager LF, Koelzer VH, Stuber R, Thoo L, Keller I, Koeck I, Langenegger M, Simillion C, Pfister SP, Faderl M et al. 2017. The ESRP1-GPR137 axis contributes to intestinal pathogenesis. eLife 6.

Mili D, Georgesse D, Kenani A. 2015. Localization and role of RAP55/LSM14 in HeLa cells: a new finding on the mitotic spindle assembly. Acta Biochim Pol 62: 613–619.

Monzo C, Haouzi D, Roman K, Assou S, Dechaud H, Hamamah S. 2012. Slow freezing and vitrification differentially modify the gene expression profile of human metaphase II oocytes. Human reproduction (Oxford, England) 27: 2160–2168.

Nilsen TW, Graveley BR. 2010. Expansion of the eukaryotic proteome by alternative splicing. Nature 463: 457–463.

Overlack K, Primorac I, Vleugel M, Krenn V, Maffini S, Hoffmann I, Kops GJ, Musacchio A. 2015. A molecular basis for the differential roles of Bub1 and BubR1 in the spindle assembly checkpoint. eLife 4: e05269.

Ramani AK, Calarco JA, Pan Q, Mavandadi S, Wang Y, Nelson AC, Lee LJ, Morris Q, Blencowe BJ, Zhen M et al. 2011. Genome-wide analysis of alternative splicing in Caenorhabditis elegans. Genome Res 21: 342–348.

Robinson JT, Thorvaldsdottir H, Winckler W, Guttman M, Lander ES, Getz G, Mesirov JP. 2011. Integrative genomics viewer. Nature biotechnology 29: 24–26.

Rohacek AM, Bebee TW, Tilton RK, Radens CM, McDermott-Roe C, Peart N, Kaur M, Zaykaner M, Cieply B, Musunuru K et al. 2017. ESRP1 Mutations Cause Hearing Loss due to Defects in Alternative Splicing that Disrupt Cochlear Development. Dev Cell 43: 318–331 e315.

Santaguida S, Tighe A, D’Alise AM, Taylor SS, Musacchio A. 2010. Dissecting the role of MPS1 in chromosome biorientation and the spindle checkpoint through the small molecule inhibitor reversine. J Cell Biol 190: 73–87.

Sha QQ, Yu JL, Guo JX, Dai XX, Jiang JC, Zhang YL, Yu C, Ji SY, Jiang Y, Zhang SY et al. 2018. CNOT6L couples the selective degradation of maternal transcripts to meiotic cell cycle progression in mouse oocyte. The EMBO journal 37.

Sha QQ, Zhang J, Fan HY. 2019. A story of birth and death: mRNA translation and clearance at the onset of maternal-to-zygotic transition in mammals†. Biol Reprod 101: 579–590.

Sha QQ, Zhu YZ, Li S, Jiang Y, Chen L, Sun XH, Shen L, Ou XH, Fan HY. 2020. Characterization of zygotic genome activation-dependent maternal mRNA clearance in mouse. Nucleic acids research 48: 879–894.

Su YQ, Sugiura K, Sun F, Pendola JK, Cox GA, Handel MA, Schimenti JC, Eppig JJ. 2012. MARF1 regulates essential oogenic processes in mice. Science (New York, NY) 335: 1496–1499.

Swartz SZ, McKay LS, Su KC, Bury L, Padeganeh A, Maddox PS, Knouse KA, Cheeseman IM. 2019. Quiescent Cells Actively Replenish CENP-A Nucleosomes to Maintain Centromere Identity and Proliferative Potential. Dev Cell 51: 35-48.e37.

Thornton BR, Toczyski DP. 2003. Securin and B-cyclin/CDK are the only essential targets of the APC. Nat Cell Biol 5: 1090–1094.

Touati SA, Buffin E, Cladiere D, Hached K, Rachez C, van Deursen JM, Wassmann K. 2015. Mouse oocytes depend on BubR1 for proper chromosome segregation but not for prophase I arrest. Nat Commun 6: 6946.

Touati SA, Wassmann K. 2016. How oocytes try to get it right: spindle checkpoint control in meiosis. Chromosoma 125: 321–335.

Vogt E, Sanhaji M, Klein W, Seidel T, Wordeman L, Eichenlaub-Ritter U. 2010. MCAK is present at centromeres, midspindle and chiasmata and involved in silencing of the spindle assembly checkpoint in mammalian oocytes. Molecular human reproduction 16: 665–684.

Wang ET, Sandberg R, Luo S, Khrebtukova I, Zhang L, Mayr C, Kingsmore SF, Schroth GP, Burge CB. 2008. Alternative isoform regulation in human tissue transcriptomes. Nature 456: 470–476.

Warzecha CC, Jiang P, Amirikian K, Dittmar KA, Lu H, Shen S, Guo W, Xing Y, Carstens RP. 2010. An ESRP-regulated splicing programme is abrogated during the epithelial-mesenchymal transition. The EMBO journal 29: 3286–3300.

Warzecha CC, Sato TK, Nabet B, Hogenesch JB, Carstens RP. 2009a. ESRP1 and ESRP2 are epithelial cell-type-specific regulators of FGFR2 splicing. Mol Cell 33: 591–601.

Warzecha CC, Shen S, Xing Y, Carstens RP. 2009b. The epithelial splicing factors ESRP1 and ESRP2 positively and negatively regulate diverse types of alternative splicing events. RNA biology 6: 546–562.

Wu W, Zong J, Wei N, Cheng J, Zhou X, Cheng Y, Chen D, Guo Q, Zhang B, Feng Y. 2018. CASH: a constructing comprehensive splice site method for detecting alternative splicing events. Brief Bioinform 19: 905–917.

Yae T, Tsuchihashi K, Ishimoto T, Motohara T, Yoshikawa M, Yoshida GJ, Wada T, Masuko T, Mogushi K, Tanaka H et al. 2012. Alternative splicing of CD44 mRNA by ESRP1 enhances lung colonization of metastatic cancer cell. Nat Commun 3: 883.

Yan X, Li F, Liang Y, Shen Y, Zhao X, Huang Q, Zhu X. 2003. Human Nudel and NudE as regulators of cytoplasmic dynein in poleward protein transport along the mitotic spindle. Molecular and cellular biology 23: 1239–1250.

Zhai R, Yuan YF, Zhao Y, Liu XM, Zhen YH, Yang FF, Wang L, Huang CZ, Cao J, Huo LJ. 2013. Bora regulates meiotic spindle assembly and cell cycle during mouse oocyte meiosis. Molecular reproduction and development 80: 474–487.

Zhang J, Zhang YL, Zhao LW, Pi SB, Zhang SY, Tong C, Fan HY. 2020. The CRL4-DCAF13 ubiquitin E3 ligase supports oocyte meiotic resumption by targeting PTEN degradation. Cellular and molecular life sciences : CMLS 77: 2181–2197.

Zhang T, Li Y, Li H, Ma XS, Ouyang YC, Hou Y, Schatten H, Sun QY. 2017. RNA-associated protein LSM family member 14 controls oocyte meiotic maturation through regulating mRNA pools. The Journal of reproduction and development 63: 383–388.

